# Cortex-wide circuits for multisensory evidence accumulation and choice formation

**DOI:** 10.64898/2025.12.23.696238

**Authors:** Gerion Nabbefeld, Sacha Abou Rachid, Irene Lenzi, Emma Cravo, Simon Musall, Björn Kampa

## Abstract

Accurate decisions in natural environments require integrating sensory information across modalities. To determine where and when multisensory signals are integrated within the cortical hierarchy, we trained mice in a visuotactile evidence-accumulation task and used widefield and two-photon calcium imaging together with optogenetic perturbations to map cortical circuits. Mice showed enhanced performance in multisensory trials that matched an additive combination of the total unisensory evidence. Cortical imaging also revealed superadditive multisensory responses in parietal and frontal regions, but visual and tactile evidence was accumulated in distinct regions, with the rostrolateral visual area (RL) being more selective for visual and the frontal medial motor cortex (MM) for tactile inputs. This modality-specific segregation persisted beyond stimulus presentation with convergence into a modality-independent choice emerging only late in the trial in the anterolateral motor cortex (ALM). Two-photon imaging confirmed that ALM neurons predominantly encoded modality-independent choices with few neurons responding to specific sensory stimuli. Optogenetic inactivation of RL and frontal cortex during the stimulus or choice period further validated their respective roles in sensory evidence accumulation and choice formation. These findings reveal a hierarchical organization in which modality-specific evidence is maintained until late in the decision process, with frontal cortex turning multisensory signals into unified choices.

## Introduction

Perception and decision-making in natural environments require the brain to integrate information across multiple sensory modalities^1,2^. The neocortex is a key structure for this process and combines inputs from different sensory systems to form a coherent representation of the external world and guide behavior^2,3^. Within the cortical hierarchy, distinct areas process sensory inputs at increasing levels of abstraction, from the encoding of basic stimulus features in primary sensory areas to the formation of modality-independent percepts and behavioral choices in higher association areas^4,5^. However, although this process is fundamental for coherent behavior, the cortical mechanisms that transform distinct modality-specific inputs into unified, sensory-independent representations remain poorly understood.

Primary unisensory areas are largely specialized for processing inputs from a single modality, but cross-modal interactions have been reported between primary visual, auditory, and somatosensory cortices in different species^6–12^. Although these effects already indicate early integration, they are typically modest and may also reflect motor activity or top-down feedback rather than genuine integration of multisensory evidence^13,14^. A much larger fraction of multisensory neurons is found in association areas located at the borders of primary sensory domains. The posterior parietal cortex (PPC) plays a particularly prominent role and has been proposed to integrate visual, tactile and auditory information^9,15–18^, but studies disagree on whether PPC itself or downstream regions in the secondary motor cortex (MOs) serve as the cortical locus for multisensory decisions^16,17,19–22^. These discrepancies are likely related to differences in task complexity: simple sensory decisions could be formed earlier in the processing hierarchy, whereas more demanding decisions that require evidence accumulation over time or modalities rely more strongly on later cortical stages^23–25^.

Multisensory integration can therefore be divided into early and late integration stages of cortical processing^1,6,26,27^. Early integration occurs at lower levels of the cortical hierarchy and involves stimulus-driven combination of sensory inputs^26,28,29^. Late integration involves higher-order cortical areas and is characterized by flexible, attention-dependent processes that adaptively weight sensory inputs based on context, expectations and task demands^1,27^. Previous studies have shown that unisensory evidence accumulation recruits distributed cortical networks operating across multiple timescales, but how these dynamics extend to multisensory integration remains unclear^6,22–25,30^. In particular, it is not known whether sensory evidence from different modalities is already combined early into a shared multisensory representation or accumulated separately within modality-specific circuits that converge only later in the decision-making process.

A compelling test case to examine how modality-specific evidence evolves along the cortical hierarchy is the integration of visual and tactile information. These modalities often complement each other during discrimination tasks, providing convergent information about object properties such as shape and texture^31–33^. Within PPC, the rostrolateral visual area (RL) is a prominent site for visuotactile integration^9^. RL is located between the primary visual (V1) and somatosensory cortex (S1) and contains a large population of neurons that respond to visual and tactile stimuli^9^. Moreover, RL has been implicated in visual evidence accumulation and decision-making and it projects to frontal cortical regions, making it an ideal candidate for early integration of visuotactile information and subsequent routing to choice-related motor regions^23,24,34,35^. This raises two central questions: does multisensory integration occur mostly locally within RL or at later stages of the cortical hierarchy, and when during the decision process are modality-specific representations transformed into unified choices? Addressing these questions requires a task that includes both sensory modalities and allows the temporal evolution of evidence integration to be tracked across cortical regions.

We therefore trained mice in a novel visuotactile evidence accumulation task and combined large-scale widefield and two-photon imaging with optogenetic perturbations to identify critical regions along the cortical hierarchy and determine if and when modality-specific information converges into a shared representation. Our results show that visual and tactile evidence are accumulated in largely segregated cortical regions, namely RL for visual and MOs for tactile inputs, while convergence to modality-independent choices occurs only later in frontal cortex. These findings suggest that the cortex maintains modality-specific representations until surprisingly late into the decision process, with full cross-modal integration arising as a final stage of evidence accumulation within frontal association areas.

## Results

### Mice additively integrate visuotactile information in a multisensory evidence accumulation task

To study multisensory perception and decision-making, we developed a novel two-alternative forced-choice task where head-fixed mice were trained to accumulate visual and/or tactile evidence over time (Figure 1a). Visual stimuli were large bars in the form of single-cycle sine waves, moving over two monitors on either the left or right side of the animal for 0.5 seconds per stimulus. Tactile stimuli were mild air puffs from two spouts in front of the animal, directed at the lower distal parts of the whiskers. In each trial, mice were presented with 3-second-long random sequences of up to six stimuli on their left and right side and had to identify the target side that contained the higher total number of stimuli (Figure 1a, right). Similar to earlier unisensory tasks, this stochastic stimulus presentation allowed us to isolate the perceptual impact of individual stimuli in the sequence on the animals’ subsequent choices^36–38^. After a 0.5 second delay, two water spouts were moved within reach of the mice which reported their decision by licking and received a water reward when correctly identifying the spout on the target side.

**Figure 1:**
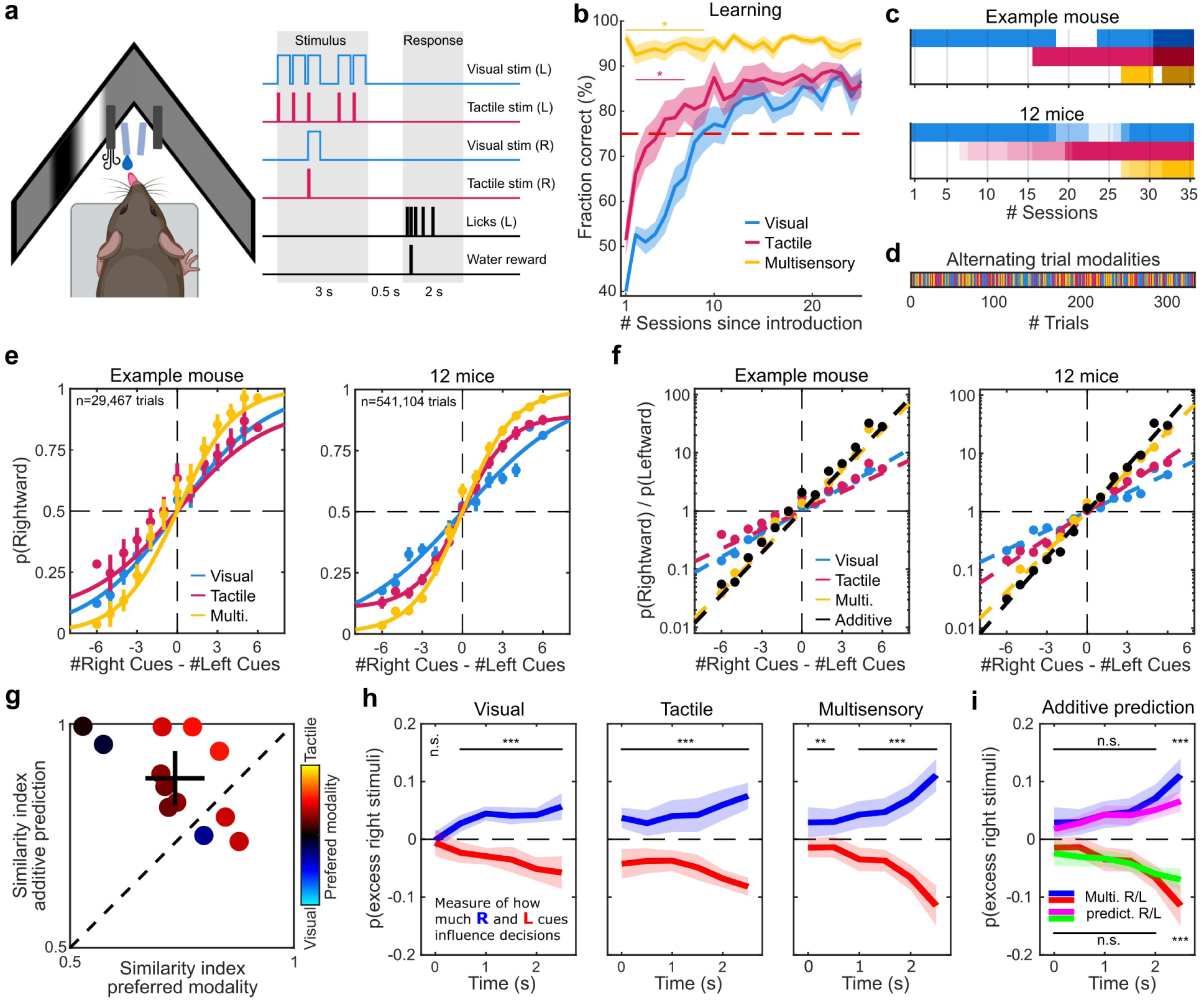
Mice additively integrate visuotactile stimuli in a multisensory evidence accumulation task. **(a)** Left: Illustration of the experimental setup. Mice were placed on a running wheel with two monitors for visual stimulation, air spouts for tactile stimulation and movable water spouts to report choices and obtain rewards. Right: Schema of an example multisensory discrimination trial with coherent visual and tactile stimulus presentation. After the delay period, mice reported their choice by licking one of two water spouts and received a water reward when identifying the high-rate target side. (**b**) Mice were sequentially trained on the three stimulus modalities using a detection paradigm. Visual detection was trained first, followed by tactile and lastly multisensory detection. Learning curves, shown relative to the introduction of each stimulus modality. Performance shown as mean ± s.e.m. across 12 mice. Learning curves were compared using a rank-sum test, assessing tactile versus visual performance (pink star) and multisensory versus the best unisensory performance (yellow star). (**c**) Top: Training regime for an example mouse, with colors indicating presentation of different stimulus modalities for each session. Darker shading indicates discrimination sessions. Bottom: Overview of the training regime across mice. Color intensity indicates the fraction of mice performing each stimulus modality. For visualization, discrimination sessions are not shown. (**d**) Example session with visual, tactile and multisensory stimulus trials. Different stimulus modalities were randomly interleaved across trials when they were shown in the same session. (**e**) Psychometric curves for visual, tactile and multisensory trials. Differences in the total number of right and left stimuli show task difficulty. Left: discrimination performance of an example mouse. Right: discrimination performance for all trials pooled across 12 mice. Markers and error bars indicate the mean performance and 95% confidence interval (CI). (**f**) Same data as in (e) but plotted as the odds ratios of right- over leftward choices on a log-scale. Black markers indicate the performance prediction when additively combining unisensory performances by adding odds-ratios from the visual and tactile condition. (**g**) Comparison of multisensory performance to the preferred unisensory condition and the additive prediction. The similarity index was computed by comparing the fitted slopes as shown in f). The cross indicates the mean and 95% CI of the two slope similarity indices. Marker color indicates the unisensory modality preference as the ratio of visual and tactile slopes. (**h**) Reverse correlation analysis, determining the impact of individual sensory stimuli on animals’ decisions. Blue line indicates the probability of observing more stimuli on the right versus left side, across all right-choice trials. Red line indicates the probability of observing more stimuli on the left versus right side, across all left-choice trials. Significant modulation was tested by comparing the right- and left stimulus probabilities in each condition (student’s t-test, n=12 mice). (**i**) Same as in h) for multisensory trials. The purple and green lines show the additive integration prediction, generated by averaging the stimulus weights in visual and tactile conditions. Differences in each time bin were tested with an LME model (n=12 mice). Significance in all panels is indicated as *, ** and *** representing p<0.05, p<0.01 and p<0.001, respectively.

We first trained mice to associate sensory stimuli and reward, by only presenting stimulus sequences with the maximum number of stimuli on the target but no stimuli on the opposing non-target side. We used this detection paradigm to first train mice with only visual and subsequently with tactile stimuli (Figure 1b). For visual detection, mice reached the criterion of 75% correct responses after 8.2 ± 1.1 days (mean ± s.e.m., n=12 mice). Once mice successfully learned the visual detection condition (at least 75% correct responses in three out of four consecutive sessions), we introduced tactile trials with 50% presentation probability. Interestingly, mice learned the tactile condition significantly faster compared to vision, relative to the first session when a given modality was introduced (4.8 ± 1.4 days, mean ± s.e.m., *p* = 0.025, ranksum test, n = 12 mice; Figure 1b). Finally, once mice reliably performed randomly alternating visual and tactile detection trials, we introduced multisensory trials with simultaneous visual and tactile stimulation (Figure 1c,d). All mice immediately performed the multisensory task condition above the expert criterion with significantly higher task performance compared to the best unisensory condition on the first day (Performance_Multisensory_ = 96.3% ± 1.3%; Performance_Unisensory_ = 90.7% ± 1.5%; mean ± sem, ranksum test: *p* = 0.01; n = 12 mice). This shows that no additional learning was required to solve the multisensory task once the animals had learned both unisensory conditions. Furthermore, the higher success rates in multisensory compared to both unisensory conditions shows that all mice could readily integrate information from both sensory modalities to improve their decisions.

Once mice achieved expert performance in the detection trials for all three randomly alternating stimulus conditions (Figure 1d), we introduced the more difficult discrimination trials by randomly drawing stimulus sequences for both the target and non-target side (presentation probabilities: 70% and 30% per stimulus and time bin for target- and non-target side, respectively^38^). All mice were able to accurately perform this discrimination task across all conditions. Comparing the performance across different difficulty levels (based on the total difference of stimuli between the target and non-target side), again showed that mice achieved significantly higher performances in multisensory trials compared to the individually preferred unisensory condition (Linear-mixed effects (LME) model to control for task difficulty and animal identify, comparing performance across modality conditions, *p* = 2.8 x 10^−4^, T = 3.7, n = 12 mice; Figure 1e). In agreement with earlier work^22^, we also found that the animals’ performance was very well described by a linear dependence on the difference in the number of left and right stimuli, suggesting that animals were summing up the information across both sensory modalities on each side to form a decision (Figure 1f).

Moreover, expressing the performance as log-odds ratios enabled us to predict the performance in multisensory trials based on the individual unisensory conditions, by assuming that stimuli in both sensory modalities were additively combined to increase the log-odds ratio of producing the correct response (Figure 1f). We also computed a similarity index comparing the slopes of the linear fits for multisensory performance to the slopes from either the preferred unisensory modality or the prediction of adding up both unisensory slopes (Figure 1g, see Methods). Indeed, we found that the animals’ performance in multisensory trials was well predicted by a purely additive combination of visual and tactile trial performances. Across all mice, the multisensory performance was best explained by additive integration of both modalities compared to the individually preferred unisensory condition (Similarity_Integration_ = 0.879 ± 0.061; Similarity_Unisensory_ = 0.735 ± 0.066; mean ± 95% CI, two-sided signrank test, *p* = 0.02, n = 12 mice, Figure 1g). Furthermore, the multisensory performance was consistently higher than the preferred unisensory performance in all mice (Supplementary Figure S1). This demonstrates that, across all mice, the multisensory performance was best explained by an additive integration of both unisensory signals rather than relying on a single unisensory modality alone.

Next, we investigated if animals also additively accumulated unisensory evidence over time. Taking advantage of our stochastic stimulation paradigm, we performed a reverse correlation analysis to determine how sensory stimuli at different timepoints during the stimulus period influenced the animals’ decisions (Figure 1h). For each modality, left- and rightward decisions were associated with a corresponding increase of excess stimuli on the target side that increased over time of stimulus presentation. Mouse choices were therefore more affected by late versus early stimuli. However, with the exception of the first visual stimulus, animals integrated sensory stimuli from the entire stimulus period (LME model, *p* = 3.7 x 10^−3^ to 7.2 x 10^−12^ across bins, n = 12 mice). Since the reverse correlations showed very similar tendencies across all modalities, we again tested if stimulus weights during multisensory trials could also be predicted by an additive integration of visual and tactile stimuli at individual time points. Indeed, the probability of excess multisensory stimuli for left and right choices was predicted by a linear combination of the unisensory stimulus probabilities with no significant difference for the first five time bins (LME model, *p* = 0.51 to 0.97 across bins, n = 12 mice; Figure 1i). The only exception was a significantly higher probability of excess multisensory stimuli during the last time bin compared to the additive unisensory prediction (LME model, *p_left_* = 4.5 x 10^−6^, *p_right_* = 1.2 x 10^−4,^ n = 12 mice).

In summary, mice showed strong performance in all three interleaved modality conditions with significantly higher multisensory performance compared to their individually preferred unisensory condition. This increase occurred immediately in the first session and was explained by an additive integration of visual and tactile stimuli, demonstrating that mice equally integrated sensory information from each modality to optimize decisions.

### Combined visuotactile stimuli evoke superadditive cortical population responses

To study the cortical representation of sensory stimuli during the visuotactile evidence accumulation task, we used widefield Ca^2+^-imaging, measuring neural activity over the entire dorsal cortical surface (Figure 2a). During correctly responded detection trials, where stimuli were only presented on one side, we found that each stimulus condition evoked distinct cortical activity patterns during the stimulus and subsequent delay period that became more similar toward the response period (Figure 2b,c). Visual stimuli primarily evoked strong activation in the primary visual cortex (V1) and higher visual areas (HVAs) during the stimulus period, especially within the rostrolateral visual area (RL). HVA activity also persisted in the delay period with increasing activity in frontal cortical areas. Although weaker in comparison, we also found consistent activation of the frontal secondary motor area (MOs) that strongly increased in the delay period. In addition to the strong activation in the contralateral hemisphere, we also found activation of the binocular region of visual areas on the ipsilateral hemisphere, likely because the full-field visual stimulus covered both binocular and monocular regions of the animals’ field of view (Figure 2d). In contrast, tactile stimuli evoked stronger responses in the whisker-related part of the primary somatosensory cortex (S1) and MOs during the stimulus period, especially the posteromedial region, matching the medial motor cortex (MM) ^39^. Activity in the MOs increased further during the delay period and became more comparable between the modalities.

**Figure 2.**
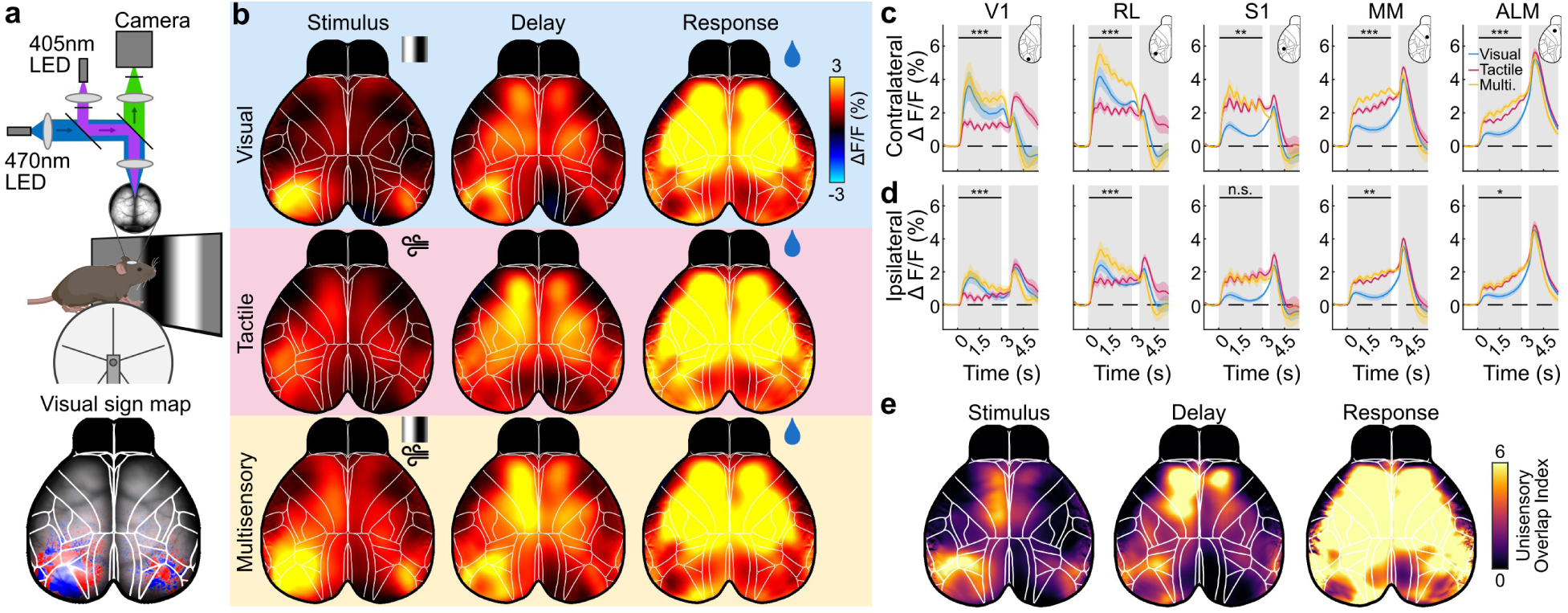
Combined Visuotactile stimuli evoke superadditive cortical population responses. **(a)** Schematic of the widefield Ca^2+^-imaging setup. Top: activity of the dorsal surface of cortex recorded under alternating illumination with blue and violet light to correct for hemodynamic signals. Bottom: Imaging data was aligned to the Allen CCF based on anatomical landmarks and confirmed using visual field sign mapping of the left visual cortex, matching the known locations of primary- and higher visual areas. (**b**) Average cortical activity of mice during different stimulus modalities and trial periods. Activity is shown for rewarded detection trials without non-target stimuli (n = 100 sessions from 4 mice). Left- and right trials were combined by reflecting data from left-target trials along the midline to represent contra- and ipsilateral activity relative to the target side instead of the left and right hemispheres, respectively. The ‘Stimulus’ column shows the average activity in the first second of the stimulus period, ‘Delay’ shows the 0.5 s delay period and ‘Response’ the first 0.5 s in the response period. (**c**) Average activity traces for different areas in the hemisphere contralateral to the stimulus. Shaded regions indicate the stimulus and response period, respectively. Insets show the cortical location for each area. Significance indicates higher multisensory activity compared to both unisensory conditions during the stimulus period, tested using an LME model. *, ** and *** represent p<0.05, p<0.01 and p<0.001, respectively. (**d**) Same as in (c) but for areas in the ipsilateral hemisphere. (**e**) Cortical maps of unisensory overlap, showing areas that were reliably activated in both visual and tactile trials. Overlap was computed as the product of normalized visual and tactile trial responses. Overlap maps are shown for the same time periods as in (b).

However, despite these modality-specific differences, cortical responses were fairly broad and most areas responded to both stimulus conditions. To isolate the areas with the largest overlap between both modalities we therefore first normalized the visual and tactile response maps for each modality and then computed their overlap as the product of both maps (unisensory overlap index, Figure 2e, Supplementary Figure S3). During the stimulus period, we found the strongest overlap in the HVAs, particularly in area RL, and the frontal area MM. The overlap in the MOs increased further during the delay period and spread throughout the cortex during the response period. These results suggest that the parietal HVAs and medial MOs are particularly well-suited for multisensory processing during the stimulus and delay period while the whole cortex became equally activated during the response period. Correspondingly, multisensory stimulation caused stronger responses throughout the cortex, with particularly strong activation of the parietal and frontal cortex (Figure 2b,c).

To test if the cortical responses to multisensory stimuli could be described by an additive combination of visual- and tactile-evoked stimulus responses, we first compared multisensory responses to the sum of unisensory responses^2^. However, multisensory responses remained below the unisensory sum across the cortex (Supplementary Figure S4). A likely reason for this result is that sensory responses only represent a fraction of the cortical activity in each trial^40^ and therefore need to be isolated from other activity patterns that co-occur in visual and tactile trials. We therefore designed a linear encoding model to fit cortical activity based on task events and movements^22,41–43^ (Figure 3a). The model used stimulus information, the general trial time and past, present and future choices and their outcomes, as predictors to explain task-related cortical activity. In addition, we used data from two behavioral cameras of the animals’ face and body, together with licking and running behavior, to predict cortical activity related to movements. All variables were then implemented as regressor sets and used jointly to predict the cortical activity of mice while performing the multisensory task. Across all discrimination sessions, the full model achieved high cross-validated explained variance (cvR^2^_full_ = 55.02% ± 0.66%; mean ± s.e.m., n = 63 sessions from 4 mice, Figure 3b), demonstrating that it was well-suited to reliably predict cortical activity and isolate sensory responses from ongoing behavior. A model based on sensory stimuli alone explained cvR^2^_stimulus_ = 15.84% ± 0.67% of the cortical variance while a movement-only model explained a larger amount of cvR^2^_movement_ = 38.46% ± 0.76%. This further confirmed the need to account for movement-related cortical responses to isolate the cortical signature of multisensory integration.

**Figure 3.**
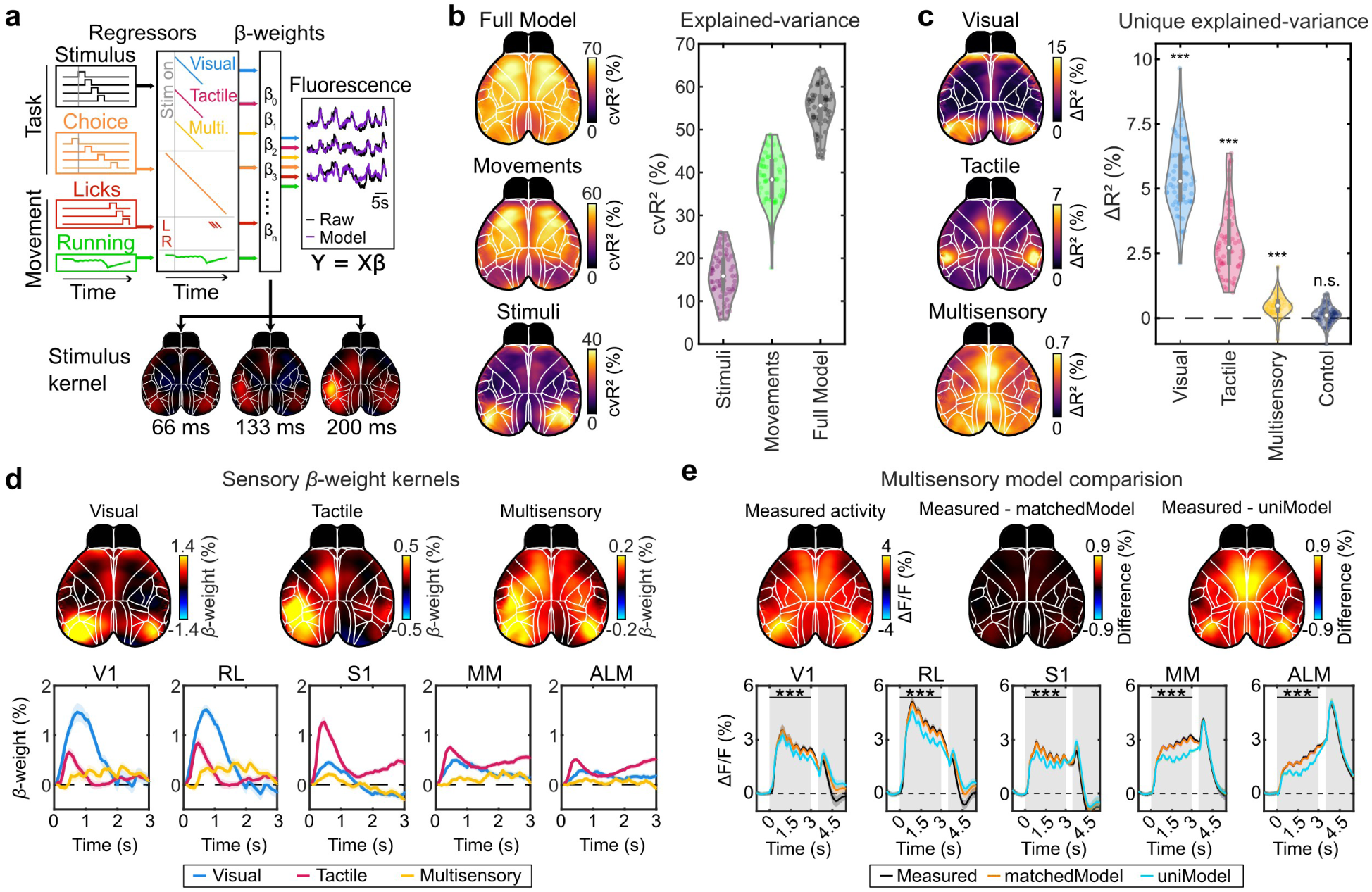
A linear encoding model reveals superadditive multisensory responses. **(a)** A linear encoding model to isolate stimulus-evoked activity from other task- and behavior-related cortical activity. The model allowed us to test if multisensory stimulus responses are fully described by the linear combination of visual and tactile responses or require an additional multisensory response variable. The model was designed to fit cortical responses to visual and tactile stimuli, independent of whether they were presented in uni- or multisensory trials. Multisensory regressors were used to provide an additional multisensory response variable to better fit cortical activity. (**b**) Predictive power of sensory stimuli and movements, compared to the full model. Cross-validated explained variance (cvR^2^) shows the amount of variance that is explained by each set of variables. Left: Spatial maps of the average cvR^2^ for the full model or a reduced movement or stimulus model. Right: cvR^2^ of sensory stimuli and movements compared to the full model over 63 sessions from 4 mice, averaged over the cortex. ( **c**) Unique explained variance (ΔR^2^) of different sensory regressors. ΔR^2^ was computed as the difference between the cvR^2^ of the full model and reduced models where a given regressor set was shuffled. Left: average spatial maps of ΔR^2^ for visual, tactile or multisensory stimuli. Right: ΔR^2^ for each stimulus, compared to a shuffle control across individual sessions, averaged over dorsal cortex. Significance was based on an LME test for the cvR^2^ of the full versus reduced models; n = 63 session from 4 mice. (**d**) Sensory β-weight kernels. Top: spatial kernels for rightward presentation of different stimulus types, averaged of the first second after stimulus onset. The β-weight sign indicates whether cortical activity increases (positive) or decreases (negative). Bottom: β-weight traces for contralateral visual (blue), tactile (pink), or multisensory (yellow) stimulation in different areas. (**e**) Comparison of a model trained solely on unisensory trials (uniModel) and a model trained on a matched number of trials from all stimulus conditions (matchedModel). Both models were validated on the same test set of multisensory trials. Top: Spatial maps of the absolute deviation between measured cortical activity (Measured, left) and predictions from the matchedModel (middle) and uniModel (right). Bottom: Averaged measured and predicted activity for the multisensory test trials in different areas. *** denotes significance for p<0.001; LME model against zero, n=66 sessions from 4 mice.

Using the encoding model we then sought to address whether multisensory responses could be predicted by a linear combination of the visual and tactile responses, or if an additional multisensory component was required to account for cortical responses when both stimulus modalities were presented simultaneously. We first isolated the variance in cortical activity that was uniquely explained by each of the stimulus conditions by generating reduced models where the visual, tactile or multisensory stimulus information was removed by shuffling the corresponding regressors in time. By comparing the explained variance of these reduced models to the full model, we obtained a conservative measure of the unique contribution of each stimulus variable to the prediction of cortical activity that could not be explained by any other model variable (unique explained variance, ΔR^2^). Visual stimuli had the largest unique contribution (ΔR^2^_Visual_ = 5.49% ± 0.18%, mean ± s.e.m., averaged over the entire dorsal cortical surface; LME test against full model, *p* = 2.7 x 10^−61^, T = 31.7, n = 63 sessions from 4 mice). Matching the averaged activity in Figure 2b, ΔR ^2^_Visual_ was highest in the occipital cortex, but also visible in the anterior and medial frontal cortex (Figure 3c). ΔR ^2^ for tactile stimuli was lower but comparable to vision (ΔR^2^_Tactile_ = 2.98% ± 0.17%, LME test against full model, *p* = 5.4 x 10^−33^, T = 16.5). ΔR^2^_Tactile_ was highest in the whisker-related part of S1 and area MM. Finally, we computed the ΔR^2^ for multisensory stimuli. The multisensory regressors were used in addition to unisensory regressors and therefore only reflected cortical activity that occurred in addition to the activity that was already predicted by the visual and tactile stimulus regressors. These additional multisensory regressors, allowing to fit non-additive multisensory responses, had smaller but still significant unique contribution in explaining cortical activity (ΔR^2^_Multisensory_ = 0.47% ± 0.05%; LME test against full model, *p* = 1.7 x 10^−6^, T = 5.0). This demonstrates that there is a significant non-additive cortical response component that cannot be explained by an additive integration of visual and tactile stimulus responses alone. In contrast to unisensory stimuli, ΔR^2^_Multisensory_ was more distributed across the cortex but showed a localized contribution in area MM, suggesting that it might have a particular role in multisensory integration.

We then investigated the full models’ β-weight kernels, representing the cortical activity assigned to the occurrence of visual, tactile, and multisensory stimuli. Since multisensory kernels were used in addition to visual and tactile kernels, this could show if multisensory stimuli lead to a superadditive increase or rather a reduction in cortical activity compared to the linear combination of visual and tactile responses. The visual and tactile kernels were largely in agreement with our earlier results, showing clear activation of V1, RL and parts of MOs while tactile kernels show stronger activity in whisker-S1 and MM (Figure 3d). Moreover, we still observed some overlap between modalities, even in primary sensory areas, with tactile responses in V1 and RL and visual responses in S1, suggesting that these regions also contain cross-modal responses that are not explained by behavior alone^13^. Interestingly, the multisensory kernel also revealed an increase in activity, especially in the HVAs and MOs (Figure 3d, top right), suggesting that multisensory stimulation induced an additional increase in cortical activity beyond the linear combination of visual and tactile responses.

However, a potential concern for this result could be that the ridge regularization of the linear model enforces an overlap of correlated β-weights which could result in part of the unisensory responses being captured by the multisensory β-weight kernels. To directly address this concern and better quantify the extent of this additional multisensory response component we therefore used an alternative approach. We first fit the model only on the unisensory trials in a given session (‘uniModel’, Figure 3e) and compared its prediction to a model that was trained on the same number of trials but also included multisensory trials in the training set (‘matchedModel’). We then validated the uni- and matched models on the same set of multisensory test trials that were excluded from the fit of both models. The only difference between the model predictions was therefore the multisensory response component that could neither be explained by the animal behavior, nor by the additive response to the combined visual and tactile stimuli. As expected, the matchedModel captured the average activity in the multisensory test trials across all cortical areas with high accuracy (Difference between measured activity and matchedModel prediction: ΔF/F_RL_ = 0.04% ± 0.02%, LME test against measured, *p* = 0.43, T = 0.79; ΔF/F_MM_ = 0.01% ± 0.01%, *p* = 0.79, T = 0.26; n = 66 sessions from 4 mice). In contrast, the uniModel significantly underestimated the average multisensory activity, especially in the parietal and medial frontal cortex during the stimulus period (Difference between measured activity and uniModel prediction: ΔF/F_RL_ = 0.47% ± 0.04%, LME model test against measured, *p* = 2.4 x 10^−13^, T = 8.17; ΔF/F_MM_ = 0.58% ± 0.03%, *p* = 3.0 x 10^−31^, T = 15.47). Together, these results demonstrate that multisensory stimulation drives stronger cortical activity as expected from a linear combination of visual and tactile responses. This effect was most visible in the parietal and frontal cortex, where the overlap of cortical responses to unisensory stimulation was also the strongest (Figure 2e). A potential consequence could be that these regions, namely areas RL and MM, are particularly involved in combining visual and tactile information to guide behavior.

### Modality-specific evidence accumulation in parietal and medial frontal cortex

To link the cortical activity to the animals’ task performance, we sought to identify activity patterns that were either related to the representation of sensory evidence or choice formation. First, we focused on the cortical representation of task-related sensory evidence in discrimination trials. Here, the target side is not directly identified by the presence of an individual sensory stimulus but requires the accumulation of evidence over time to identify the target side with the higher total number of stimuli. To identify which cortical regions reliably represented this accumulated sensory evidence, we computed the area under the receiver operating characteristic curve (AUC) for the high-rate target versus the non-target side. The AUC was computed by comparing the activity between the two cortical hemispheres for each pixel, with AUCs above 0.5 indicating higher activity in the hemisphere that was contralateral to the target side. Moreover, the AUC was computed for a balanced amount of correct and incorrect choice trials, to ensure AUCs only represented sensory evidence for the target side independent of animal choices^39^. We computed these evidence-related AUCs separately for visual, tactile and multisensory trials (Figure 4).

**Figure 4.**
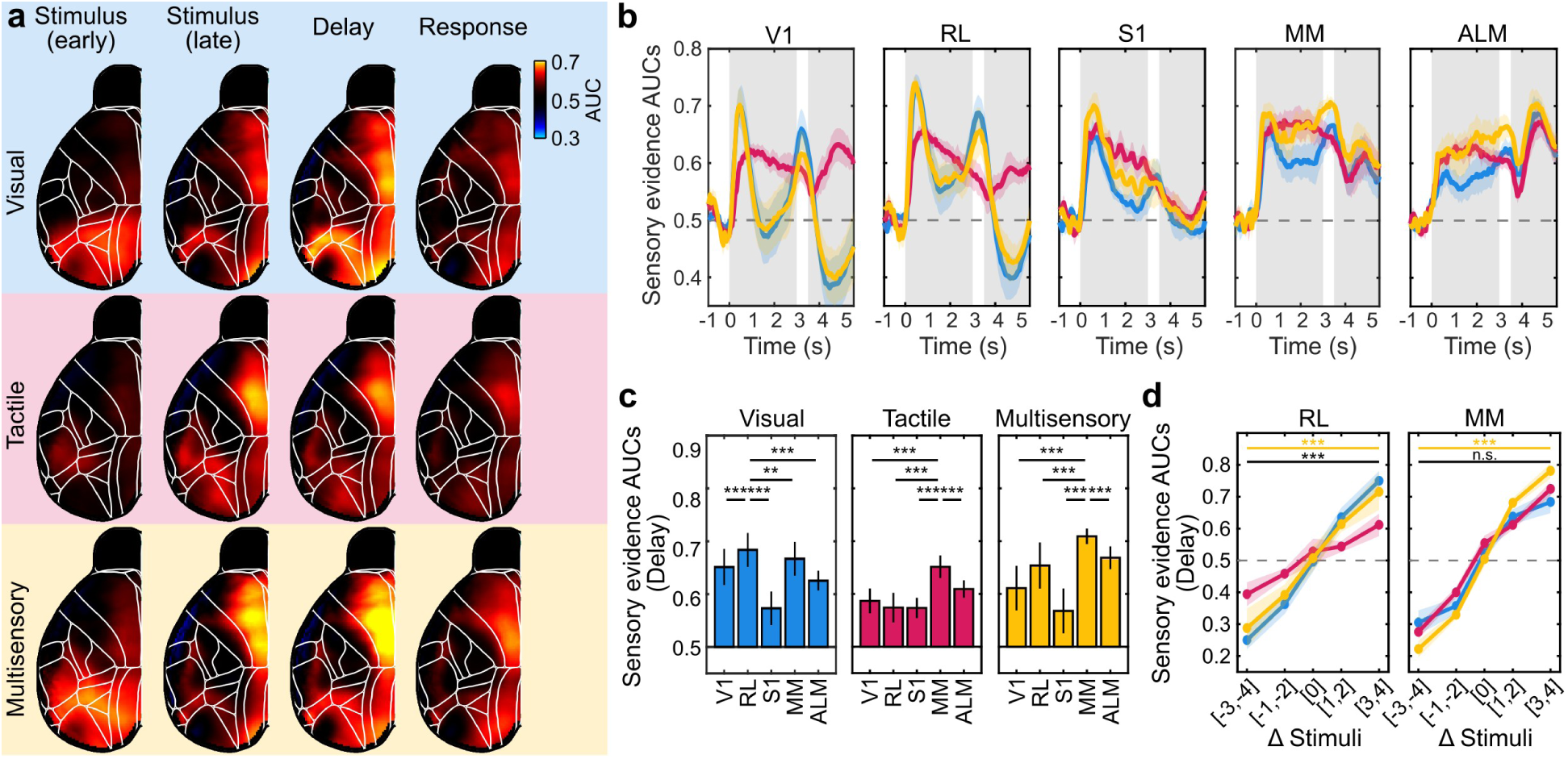
Modality-specific sensory evidence accumulation. **(a)** Representation of sensory evidence in different stimulus conditions. Presented for 0.5s time bins during the first stimulus (early), last stimulus (late), delay and response period. The AUC maps show cortical regions that reliably represented the side where the higher number of sensory stimuli was presented, based on the difference in activity between hemispheres. AUCs were computed for discrimination trials with an equal number of correct and incorrect trials to avoid an influence of choice-related activity. AUCs above 0.5 indicate a reliable representation of contralateral sensory evidence. (**b**) AUC traces for each stimulus condition in selected cortical areas. AUCs for individual mice were computed using a bootstrapping procedure to correct for different trial-counts across sessions. Traces show the mean ± s.e.m. across mice. (**c**) Average sensory AUCs during the delay period for the areas shown in (b). Bars show AUCs as mean ± s.e.m. across mice. For each stimulus condition, the area with the highest AUC was determined and tested against the other areas using an LME model. (**d**) Neurometric curves of RL and MM shown as mean ± s.e.m. across 4 mice. AUCs computed for choice-balanced discrimination trials based on the difference in stimuli on the target and distractor side (Δ Stimuli). Results are shown for easy (Δ Stimuli = [3,4]) and hard discrimination trials (Δ Stimuli = [1,2]). Black significance stars for comparison of visual and tactile AUCs; Yellow stars for comparison of preferred unisensory AUCs and multisensory AUCs. n for all panels = 4 mice, 100 bootstrap samples each. ***: p<0.001, **: p<0.01 and *: p<0.05.

In line with our earlier results, sensory AUCs were particularly high in the parietal and medial frontal cortex, with visual evidence peaking in the parietal area RL and tactile information in the frontal area MM (Figure 4a, Supplementary Figure S5). These modality-specific differences also persisted in the delay period after the stimulus sequences were over. However, while RL remained mostly predictive for the high-rate target side in visual trials, MOs became more evenly predictive for both visual and tactile trials (Figure 4a,b).

A similar effect was also visible for multisensory trials which showed equally high AUCs in the same regions as for visual and tactile trials but the most consistent target side representation was found in area MM during the delay period. The temporal dynamics of the AUCs also differed between modalities. Tactile AUCs in MM increased progressively during the stimulus presentation, whereas visual AUCs in V1 and RL sharply rose with the onset of the stimulus but decreased as the stimulus progressed. In V1, AUCs dropped to chance levels; in contrast, RL maintained elevated AUCs that peaked toward the end of the stimulus period and during the delay phase. This suggests that the initial sensory response can also indicate the target side, likely because the first stimulus in the sequence also elicited the strongest visual response difference between the contralateral and ipsilateral hemispheres (Figure 2c). However, only RL showed a sustained representation of sensory evidence that persisted into the delay period, suggesting a temporal accumulation of sensory evidence.

To compare AUCs between areas while accounting for different trial-counts across sessions and sensory modalities, we used a bootstrapping procedure. For each modality, 40 choice-balanced trials were randomly resampled with replacement from all sessions and the AUC was computed. This was repeated 100 times for each mouse to generate a bootstrap distribution of AUCs for comparison. During the delay period, RL displayed the highest visual AUCs compared to all other areas (Visual: AUC_V1_ = 0.65 ± 0.03, AUC_RL_ = 0.69 ± 0.03, AUC_S1_ = 0.57 ± 0.03, AUC_MM_ = 0.67 ± 0.03, AUC_ALM_ = 0.63 ± 0.02; LME model, *p* = 6.5 x 10^−3^ to 7.17 x 10^−69^, mean ± s.e.m., Figure 4c), indicating that RL was the most reliable accumulator of visual evidence. In contrast, tactile AUCs during the delay period were the highest in MM (Tactile: AUC_V1_ = 0.58 ± 0.02, AUC_RL_ = 0.57 ± 0.03, AUC_S1_ = 0.57 ± 0.02, AUC_MM_ = 0.65 ± 0.02, AUC_ALM_ = 0.61 ± 0.02; LME model, *p* = 2.9 x 10^−12^ to 1.2 x 10^−34^). MM also showed the highest multisensory AUCs compared to all other areas (Multi.: AUC_V1_ = 0.61 ± 0.04, AUC_RL_ = 0.65 ± 0.04, AUC_S1_ = 0.57 ± 0.04, AUC_MM_ = 0.71 ± 0.02, AUC_ALM_ = 0.67 ± 0.02; LME model, *p* = 1.4 x 10^−12^ to 3.7 x 10^−93^).

Next, we compared how RL and MM represented the target-side across different task difficulties. We used the same bootstrapping procedure but computed AUCs for easy (ΔStimuli=[3,4]) and hard (ΔStimuli=[1,2]) trials separately. AUCs in both areas scaled with the sensory evidence (Figure 4d). However, RL represented visual evidence much more reliably than tactile evidence and showed no multisensory enhancement (Visual vs. Tactile: *p*=3.7 x 10^−278^, T=-39.49; preferred Unisensory vs. Multisensory: *p*=9.1 x 10^−49^, T=-14.93; LME model). In contrast, MM represented both visual and tactile evidence equally well but encoded multisensory evidence more strongly (Visual versus tactile: *p*=0.85, T=0.19; preferred unisensory versus multisensory: *p*=6.8 x 10^−8^, T=5.41; LME model). These results suggest that the temporal accumulation of unisensory evidence from visual and tactile stimulation occurs in distinct cortical areas, specifically area RL for visual and MM for tactile information. Both regions also accumulated multisensory information but MM represented the target side most reliably in multisensory trials with visual and tactile AUCs converging during the delay period. MM may therefore play a particularly prominent role in tactile evidence accumulation and multisensory integration.

### Cross-modal choice formation in frontal cortex

Having found regional differences in the representation of accumulated visual and tactile sensory evidence, we wondered if these areas also displayed choice-related activity in a modality-specific manner. To address this question, we first used a logistic regression decoder to predict animal choices based on the entire cortical activity across all trials and modalities. This provided an overview of the time course of choice formation and how accurate choices could be predicted from cortical activity. Similar to our earlier analysis, we balanced the amount of correct and incorrect choice trials to separate choice- from stimulus-related activity patterns. The decoder reliably predicted animal choices above chance throughout the stimulus period but remained roughly at the same level (decoder accuracy_Stimulus_ = 58.3% ± 0.4%, mean ± s.e.m., n = 4 mice, Figure 5a). This was in line with our behavioral results, showing that choices were affected by sensory evidence throughout the stimulus period (Figure 1h). The prediction accuracy then sharply increased toward the end of the stimulus and over the delay period (decoder accuracy_Delay_ = 74.0 ± 1.2 %, mean ± s.e.m., n = 4 mice).

**Figure 5.**
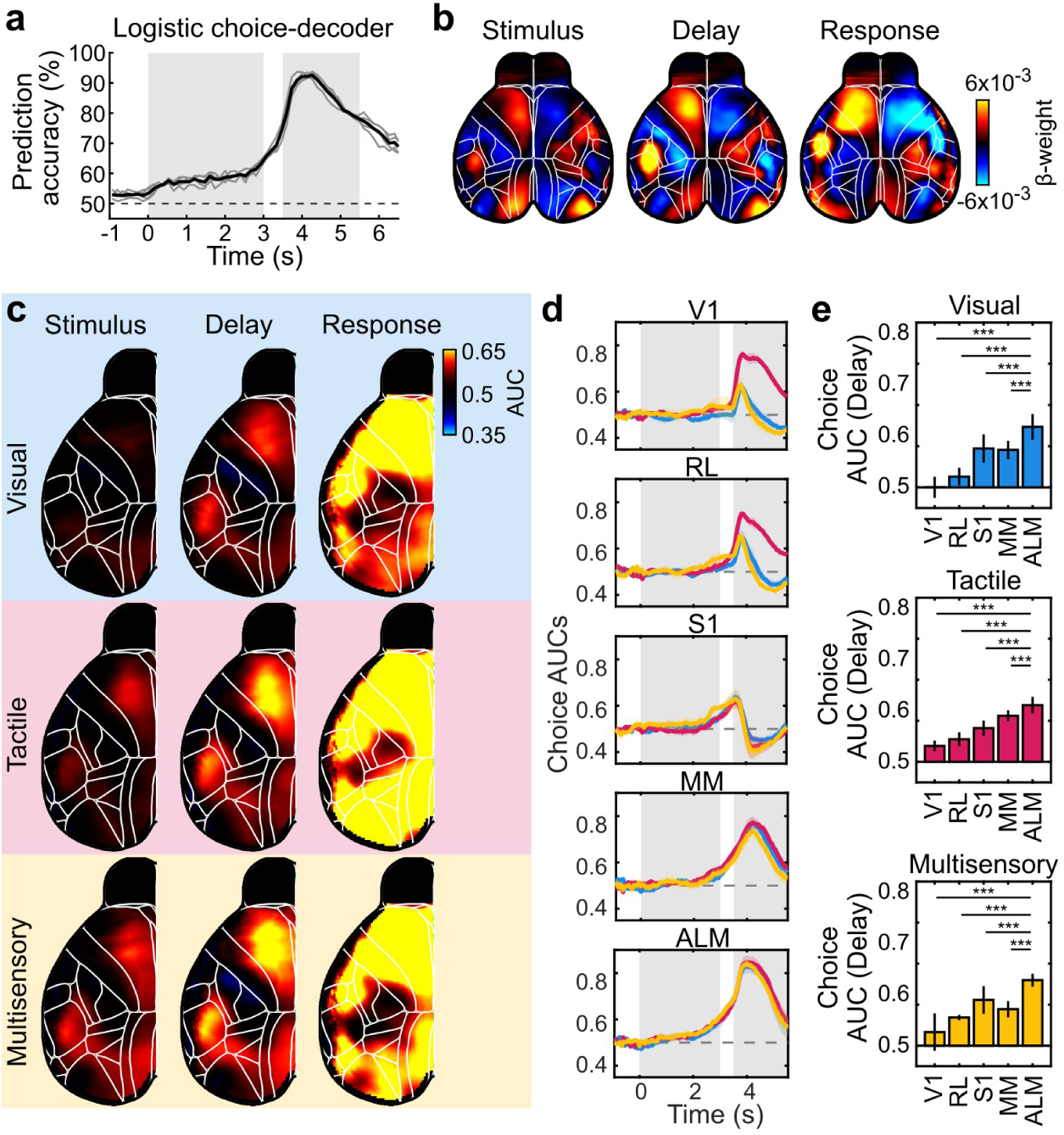
Cross-modal choice formation across cortex. **(a)** Cross-validated choice decoder accuracy, based on dorsal cortical activity to predict upcoming animal choices across different trial times. The decoder was fit separately at each time point using choice-balanced trials and 10-fold cross validation. Prediction accuracy is shown for individual mice (gray) and the mean across mice (black). (**b**) Mean decoder weights predicting choices during the stimulus (last 0.5 seconds), the delay (0.5 seconds), and the response period (first 0.5 seconds). (**c**) Spatial maps of choice AUCs during the same time-points as in b), reflecting how reliably interhemispheric activity differences predicted upcoming choices. (**d**) Choice AUC traces for each stimulus condition in selected cortical areas. Traces show the mean ± s.e.m. across mice, n = 4 mice, 100 bootstrap samples each. ( **e**) Average choice AUCs during the delay period for the areas shown in d). Bars show choice AUCs as mean ± s.e.m. across 4 mice. For each stimulus condition, the area with the highest AUC was determined and tested against the other areas using an LME model. ***: p<0.001.

Matching earlier work from an auditory task^41^, the decoder β-weights were increased across the anterior cortex with particularly large contributions from MOs and subregions of somatosensory cortex (Figure 5b). The weights were generally higher in the delay versus the stimulus period but showed the same cortex-wide patterns, suggesting that choice formation does not include different cortical areas during different trial periods. Moreover, we consistently found the same β-weight maps across visual, tactile and multisensory discrimination trials (Supplementary Figure S6), suggesting that, contrary to sensory evidence accumulation, choice formation across cortical areas is largely modality-independent.

Because the choice decoder relied on cortex-wide activity patterns, we asked whether the β-weights accurately reflected choice-related activity in all areas, particularly when the same information was represented across multiple regions. Moreover, the limited number of choice-balanced trials per stimulus condition may have obscured more subtle differences across stimulus conditions. We therefore computed AUC values for each pixel, this time to separate contra- versus ipsilateral choice trials, independent of the sensory evidence. In agreement with our earlier analysis, cortex-wide choice AUCs remained low after stimulus onset and only increased toward the end of the stimulus and during the delay period (Figure 5c,d, Supplementary Figure S7). Choice AUCs were highest in MOs, particularly in the anterolateral motor cortex (ALM) and interestingly, also in the somatosensory cortex. In contrast, most cortical regions reliably represented the animal’s choices during the response period. Therefore, we focused on activity during the delay period of the task, to determine if there are modality-specific differences in the choice formation. Across all stimulus modalities, ALM consistently showed the most reliable choice representations during the delay period (ALM: AUC_Visual_ = 0.65 ± 0.03, AUC_Tactile_ = 0.64 ± 0.02, AUC_Multisensory_ = 0.66 ± 0.02, mean ± s.e.m.; *p* = 2.7 x 10^−5^ to 3.4 x 10^−90^, LME model; 4 mice with 100 bootstrapped samples each; Figure 5e).

Together, these results suggest that choice formation occurs equally in the same frontal regions across different stimulus modalities, especially during the delay period. ALM appears to be the central cortical hub that forms decisions across sensory modalities, likely receiving accumulated visual evidence from RL and tactile evidence from MM.

### ALM neurons encode choices across modalities

Our widefield imaging results suggest that choice formation occurs in the same cortical areas, particularly in ALM, regardless of the stimulus modality. However, stimulus-specific choices could still be embedded in the same areas by recruiting different subsets of neurons. We therefore used two-photon microscopy to measure single-cell activity in the ALM of task-performing mice to test if the stimulus or choice tuning of individual neurons differs across visual, tactile and multisensory trials (Figure 6a). While ALM contained neurons responding to sensory stimuli (Figure 6b, top), the vast majority of tuned cells represented the upcoming choices (Figure 6b, bottom). We quantified the fraction of cells that responded to sensory stimuli, reliably represented the target side or the upcoming animal choices. First, we tested if a significant fraction of cells responded to sensory stimuli during the stimulus period of simple detection trials, compared to a shuffle control. The fraction of stimulus-responsive cells was generally low (∼6%) and only significantly differed from the shuffle control for tactile and multisensory but not visual stimuli (fraction _Visual_ = 5.60%, [4.56%, 6.86%], fraction_Tactile_ = 6.83%, [5.67%, 8.19%], fraction_Multi._ = 6.31%, [5.21%, 7.63%], mean, [95% CI]; *p*_Visual_ = 0.22, *p*_Tactile_ = 0.003, *p*_Multi._ = 0.02, Binomial test compared to shuffle control, n = 1553 cells from 2 mice). Next, we determined if cells reliably represented the target-side with the higher number of sensory stimuli. For this, we followed the same choice-balancing procedure previously described, comparing the activity during the delay period of discrimination trials in each modality. However, the fraction of target-selective cells was not different from the shuffle control in any condition (fraction_Visual_ = 4.38%, [3.47%, 5.51%], fraction_Tactile_ = 5.09%, [4.10%, 6.29%], fraction_Multi._ = 3.80%, [2.96%, 4.87%], mean, [95% CI]; *p*_Visual_ = 1, *p*_Tactile_ = 0.5, *p*_Multi._ = 0.85, Binomial test compared to shuffle control, n=1553 cells from 2 mice). Lastly, we determined the fraction of choice-selective cells during the delay period. Here, a larger fraction of cells reliably represented upcoming choices in all stimulus modalities (fraction_Visual_ = 9.53%, [8.17%, 11.09%], fraction_Tactile_ = 7.53%, [6.32%, 8.95%], fraction_Multi._ = 9.53%, [8.17%, 11.09%], mean, [95% CI]; *p*_Visual_ = 2.7 x 10^−8^, *p*_Tactile_ = 3.9 x 10^−4^, *p*_Multi._ = 6.1 x 10^−6^, Binomial test compared to shuffle control, n=1553 cells from 2 mice). These results show that stimulus information is only weakly represented in individual ALM neurons whereas a considerable fraction of cells represented the upcoming choices across modalities.

**Figure 6.**
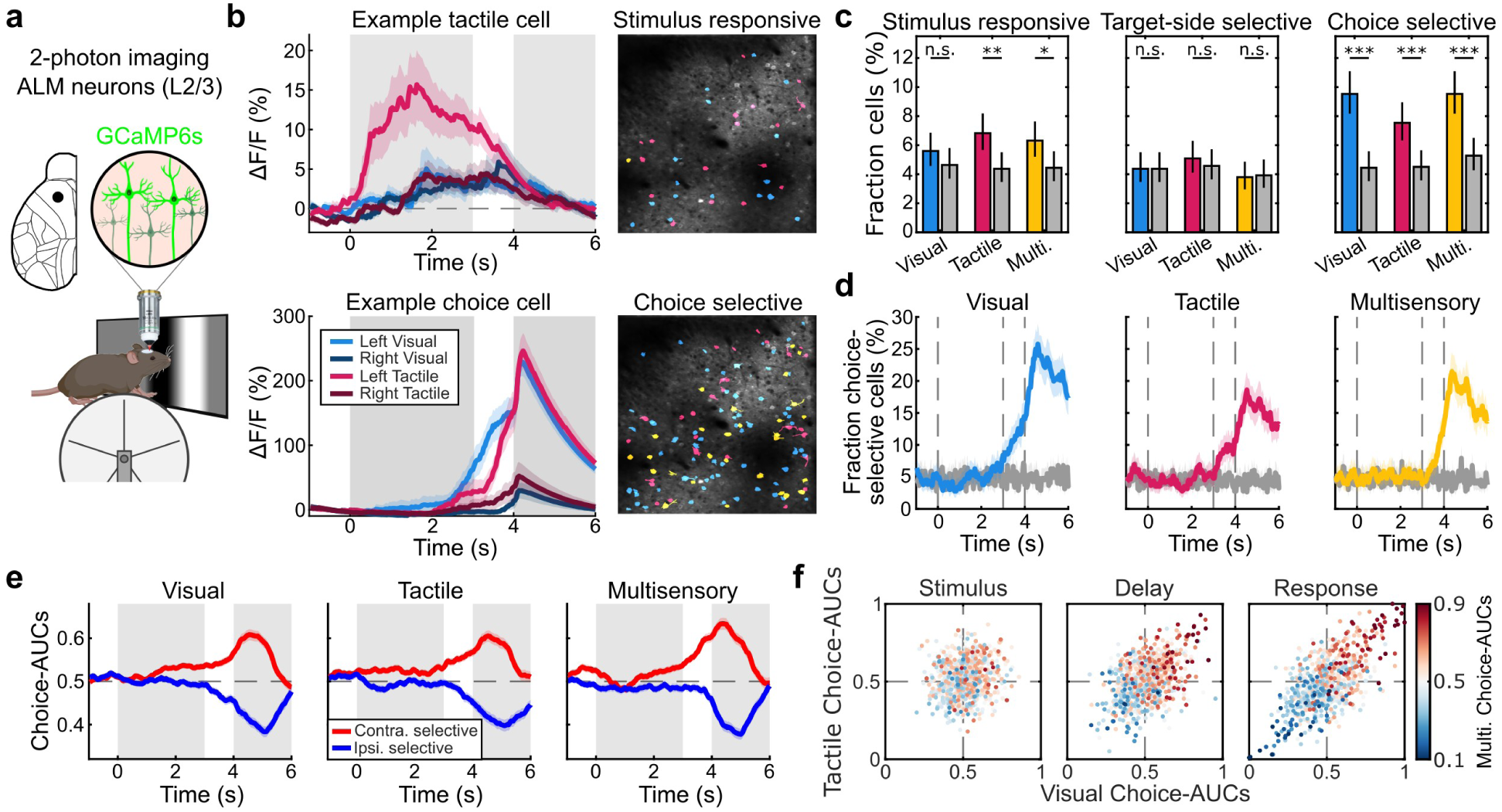
ALM neurons encode choices across sensory modalities. **(a)** Two-photon microscopy to record single cell activity in task-performing mice. (**b**) Left: Activity of example neurons averaged across visual and tactile trials with stimuli on either the left or the right side. Traces show the mean and shading the s.e.m. for a tactile-responding (top) and a choice-selective cell (bottom). Gray periods indicate the stimulus and response periods. To better characterize single-cell activity during the delay period, we extended this period to 1 second for the two-photon recordings. Right: Example field of views with stimulus-responsive (top) and choice-selective cells (bottom). Colored cells showed significant responses in visual (blue), tactile (pink) or both trial modalities (yellow). Stimulus responders were defined by a significantly higher response during the stimulus period, compared to baseline (one-sided signrank test). Choice selectivity was defined as a significant difference in activity during the delay period of choice-balanced discrimination trials (two-sided ranksum test). (**c**) Fraction of significantly task modulated cells. Left: Fractions of sensory-responsive cells during the stimulus period across modalities. Middle: Fractions of target side-selective neurons during the delay period. Right: Fraction of choice-selective neurons during the delay period. Errors indicate 95% confidence-intervals (Cis). Gray bars indicate shuffle controls by shuffling labels across trials. Binomial test, ***: *p*<0.001, **: *p*<0.01, n=1553 cells. (**d**) Fraction of choice-selective cells across time. Traces and shading indicate the mean ± CIs over the course of visual (left), tactile (middle) and multisensory (right) discrimination trials. Gray traces and shading indicate the shuffle control. Dashed lines indicate the stimulus and response period. (**e**) Choice AUCs for all choice-modulated neurons. Traces and shading indicate the mean AUC ± s.e.m. across trials for the different modalities. (**f**) Correlation of neuronal choice selectivity in visual, tactile and multisensory discrimination trials. Marker colors indicate the choice selectivity in multisensory trials. Results are shown for the last second of the stimulus period (Stimulus), the delay period (Delay), and the first second of the response period (Response).

In agreement with our widefield imaging results (Figure 5a), the fraction of choice-selective neurons increased over the course of the trial, especially in the delay and subsequent response period (Figure 6d). The increase in selective neurons was largely similar across modalities with a significant fraction of selective neurons occurring around the start time of the delay period (t_Visual_ = 3.23s, t_Tactile_ = 3.30s, t_Multi._ = 3.67s, *p*_Visual_=7.9 x 10^−3^ to 3.7 x 10^−37^, *p*_Tactile_=5.7 x 10^−3^ to 5.0×10^−25^, *p*_Multi_=1.2 x 10^−2^ to 2.6 x 10^−30^, Binomial test against shuffle control). Interestingly, reliable choice representations occurred a bit later in multisensory versus unisensory trials, potentially representing a more complex integration process despite the higher choice accuracy in multisensory trials.

To test for differences in the representation of ipsi- and contralateral choices between modalities, we next selected all cells that displayed a significant choice modulation for at least 1 second in any of the modality conditions and classified them as either ipsi- or contralateral choice selective. The fractions of ipsi- and contralaterally choice selective cells was similar across conditions (Contralateral fraction_Visual_ = 57.8%, fraction_Tactile_ = 55.0%, fraction_Multi._ = 53.6%, n=360 choice selective cells from 2 mice) and also showed a comparable time course of the average choice AUCs across the trial (Figure 6e). However, choice AUCs already increased earlier than the fraction of selective neurons, with significant contralateral tuning during the middle of the stimulus period. In contrast, ipsilateral tuning occurred much later in time, especially for the multisensory condition (Visual: t_Contra_ = 1.27s, t_Ipsi_ = 3.37s; Tactile: t_Contra_ = 2.53s, t_Ipsi_ = 3.53s; Multisensory: t_Contra_ = 1.97s, t_Ipsi_ = 3.80s).

Lastly, we asked if separate neural populations encoded choices in visual and tactile trials, or if the same neurons were equally choice-selective regardless of the stimulus modality. We therefore compared the choice-AUCs of all selective cells during the stimulus, delay and response period across modalities (Figure 6f). During the late stimulus phase, visual and tactile choice AUCs were uncorrelated (Spearman *r* = 0.14, *p* = 0.067, n = 1553 cells) and only weakly correlated between multisensory and the most-selective unisensory modality (*r* = 0.24, *p* = 0.001). In contrast, correlations strongly increased during the delay period (Visual versus tactile: *r* = 0.52, *p* = 2.3 x 10^−18^, multi- versus unisensory: *r* = 0.64, *p* = 5.9 x 10^−30^), indicating that choice tuning became more modality-independent after stimulus offset. Correlations further increased during the response period (Visual versus tactile: *r* = 0.65, *p* = 2.1 x 10^−40^, multi- versus unisensory: *r* = 0.71, *p* = 2.7 x 10^−51^). Moreover, neurons that were choice-selective in visual and tactile trials were significantly more selective compared to neurons that were only selective in one stimulus modality (bimodal choice AUC = 0.78 ± 0.006, n = 93 cells, unimodal choice AUC = 0.72 ± 0.003, mean ± s.e.m., n = 360 cells; Mann-Whitney U test, *p* = 1.6*10^−16^; Supplementary Figure S8). Together, these results suggest a convergence of visual and tactile choice information onto the same neuronal population in ALM during multisensory decision-making.

### Cortical inactivation reveals area-specific evidence accumulation and shared choice formation

Our imaging results suggest that visual and tactile evidence are preferentially accumulated in separate cortical areas whereas behavioral choices are formed in the same areas, particularly in frontal cortex. To causally test this hypothesis, we performed optogenetic inactivation of either RL or MOs in task-performing mice (Figure 7a,b). Both regions were bilaterally injected with an anterograde adeno-associated virus to express the blue light-sensitive inhibitory opsin stGtACR2^44^ in all excitatory neurons (Figure 7c). By using blue-light illumination of RL or MOs, we then bilaterally inactivated either area during the stimulus or the delay/response period to test their respective causal contribution to sensory perception and choice formation (Figure 7b).

**Figure 7.**
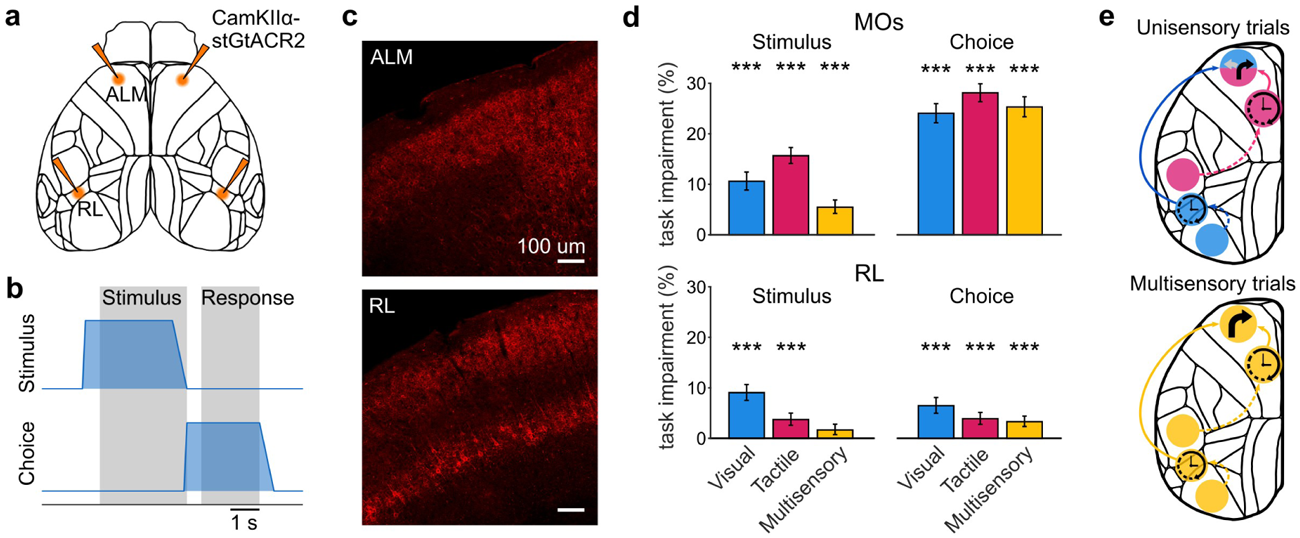
Optogenetic inactivations reveal area-specific evidence accumulation and shared choice formation. **(a)** Top: Schematic of bilateral optogenetic inactivations. Excitatory neurons in MOs and RL were transfected with a viral vector to express the soma-targeted, inhibitory opsin stGtACR2. Bottom: Schematic of bilateral inactivations during the stimulus and choice periods. **(b)** Histological validation of opsin expression in excitatory neurons of RL and MOs. **(c)** Impact of optogenetic inactivation on task performance in each modality. Task impairments are shown as the performance difference between optogenetic versus control trials across detection and discrimination trials. Stars show significant differences in performance using a binomial test (***: p<0.001, n=1569-2665 optogenetic trials from 4 mice). (**e**) Proposed circuit model of visuotactile evidence accumulation. V1 and S1 encode current sensory stimuli on a short timescales, while RL and MM accumulate sensory evidence over longer time-scales in a modality-specific manner. In contrast, ALM forms choices across stimulus modalities, indicative of modality-independent decision-making (top). In multisensory trials, visual and tactile evidence converges in ALM and increases decision reliability (bottom).

Bilateral inactivation of RL during either the stimulus or the choice period primarily impaired task performance in visual trials (Stimulus: impairment_Visual_ = 9.04% [7.52%, 10.64%], *p*_Visual_ = 9.25 x 10^−18^, impairment_Tactile_ = 3.70% [2.56%, 4.98%], *p*_Tactile_ = 1.70 x 10^−5^, impairment_Multi_ = 1.64% [0.72%, 2.79%], *p*_Multi_ = 0.12; Choice: impairment_Visual_ = 6.46% [4.95%, 8.07%], impairment_Tactile_ = 3.87% [2.75%, 5.12%], impairment_Multi_ = 3.29% [2.33%, 4.41%], *p* = 7.72 x 10^−9^ to 3.07 x 10^−6^; Binomial test compared to control trials; Figure 7d, bottom). This was in agreement with our imaging results and shows that, despite responding to both visual and tactile stimuli, RL was only causally required for visual evidence accumulation. In contrast, MOs inactivation during the stimulus period impaired performance in visual and tactile trials but with a stronger effect in tactile trials (Stimulus: impairment _Visual_ = 10.59% [8.85%, 12.42%], impairment_Tactile_ = 15.65% [14.11%, 17.27%], impairment_Multi_ = 5.48% [4.24%, 6.88%], *p* = 3.08 x 10^−75^ to 6.47 x 10^−14^). This suggests that tactile evidence is preferentially accumulated in MOs (Figure 7d, top). Furthermore, inactivating MOs during the delay and response period strongly impaired choice accuracy across modalities (Choice: impairment_Visual_ = 24.05% [22.18%, 25.95%], impairment_Tactile_ = 28.10% [26.34%, 29.90%], impairment_Multi_ = 25.30% [23.37%, 27.32%], *p* = 3.18 x 10^−212^ to 7.31 x 10^−98^). Importantly, these effects were only due to a decrease in response accuracy rather than a change in response probability ( *p* = 0.48, T = -0.71, LME model, n = 144 sessions from 4 mice; Supplementary Figure S9).

Together, our imaging and optogenetic results suggest a cortical circuit model for multisensory decision making where visual evidence is accumulated in RL and tactile evidence is accumulated in MOs, particularly in area MM (Figure 7e). During the delay period, visual and tactile information then converge in MOs with ALM emerging as a central hub for integrating multisensory signals into unified choice representations.

## Discussion

We developed a new multisensory evidence accumulation task for head-fixed mice and found that animals readily combined visual and tactile sensory information to improve their discrimination performance. Multisensory performance was best explained by an additive integration of unisensory inputs, although a linear encoding model also revealed super-additive multisensory cortical responses. These multisensory responses were most pronounced in the parietal area RL and frontal area MM. Further analysis of task-related sensory activity highlighted RL and MM as key areas for visual and tactile evidence accumulation, respectively. While unisensory evidence was integrated in largely separate cortical regions, it then converged in MOs during a cross-modal decision-making process that was reflected in the modality-independent choice tuning of individual ALM neurons. Optogenetic inactivation further confirmed this cortex-wide functional organization, revealing modality-specific impairments in RL and MOs during evidence accumulation and modality-independent effects in MOs during choice formation.

The design of our multisensory task allowed us to test if mice equally accumulated visual and tactile information and how multisensory decisions were shaped by each unisensory stimulus component. In line with earlier unisensory studies^45–47^, we found that choices were affected by all stimuli throughout the stimulus period, demonstrating that mice actively accumulated sensory evidence over time. Moreover, the behavioral impact of multisensory stimuli was well predicted by the sum of visual and tactile stimuli, suggesting that they equally attended both unisensory modalities. Correspondingly, we found an immediate and consistent enhancement in multisensory versus unisensory discrimination performance, matching a large body of literature across species on the perceptual benefits of multisensory integration^12,48–53^. Similar to a recent study showing that mice additively combine audio-visual information to identify a stimulus location^22^, our results demonstrate that integration of visuo-tactile evidence is also additive. Importantly, this integration occurred in a task requiring temporal evidence accumulation and working memory, extending previous findings to a more complex decision-making context.

To study the cortical representation of multisensory decision-making we used cortex-wide Ca^2+^-imaging. In particular, we asked if mice immediately integrate visual and tactile information to accumulate multisensory information in associative areas, such as MOs^21,22,54^ or RL^9,18,55^, or rather utilize parallel modality-specific pathways that converge at a larger processing stage to guide behavior^18,56,57^. Both RL and MOs responded to visual and tactile stimulation and showed increased responses to multisensory stimuli. Multisensory responses in both regions were superadditive, suggesting the presence of specialized circuits for spatiotemporal coincidence detection of visuo-tactile inputs. However, since the behavioral performance was best explained by an additive integration of unisensory evidence, these additional cortical responses did not translate into further improved behavioral performance^48^.

We also found a clear unisensory response bias in both regions, with RL responding preferably to visual stimulation whereas area MM, was more responsive to tactile simulation. Notably, neural encoding of tactile information in MM has also been found in a spatial object localization task^39^ and matches the same cortical region that has been referred to as whisker M1, with rapid responses to whisker stimulation and direct projections from the barrel cortex^58–60^. Beyond its role for multisensory integration and choice formation, area MM therefore appears to play an important role for tactile evidence accumulation. This was also confirmed by our optogenetic inactivation of MOs, showing a stronger impairment in discrimination performance in tactile versus visual or multisensory trials.

The visual response preference in RL aligns with its role as a higher visual area, with retinotopic responses and functional tuning to complex visual stimuli^57,61^. The prolonged representation of visual evidence into the delay period suggests that RL maintains visual evidence on longer timescales than V1, making it well-suited for visual evidence accumulation. Several studies have also implicated RL in visuotactile integration^9,18,55^, yet we found no significant impact of RL inactivation on multisensory task performance. Since RL is prominently involved in the processing of motion within the dorsal visual stream^62^ this could have favored the discrimination of the moving visual stimuli over the pulsatile tactile stimuli in our task. A recent study also found that lateral RL is activated during active whisking and causally required for the discrimination of textures^18^. Due to its connections to motor areas, such as motor cortex^63^ and superior colliculus^64,65^, and tuning preference for high temporal frequencies^57^, multisensory processing in RL might therefore be focused on active behaviors which require fast sensorimotor integration and processing of fine spatiotemporal details, such as texture discrimination, spatial navigation or prey capture^66^. RL also supports rapid cross-modal generalization in multisensory object detection when visual and tactile stimuli are closely aligned within the peri-personal space^55^, again highlighting its role in precise, spatially constrained multisensory integration. In contrast, our task required discrimination of sensory inputs from the opposite hemifields. Although we also observed a reduction in training time when introducing the second modality, all mice still required several sessions to achieve expert tactile task performance, arguing against an immediate transfer of task knowledge from one modality to another. Together, our results therefore suggest that multisensory processing in RL might be optimized for integrating sensory inputs within the same region of the sensory field but not full-field cross-hemispheric comparison, as required in our task.

For comparing multisensory evidence across hemispheres to make bilateral left-right choices, our results instead highlight the importance of MOs. This is likely due to its strong inter-hemispheric and corticothalamic feedback loops that promote the formation of side-specific choice signals over longer timescales^39,67–69^. Using a choice decoder or a pixel-wise AUC approach, we found largely similar choice-related activity in visual, tactile and multisensory trials, suggesting that choice formation in MOs is largely modality-independent. Moreover, we also found choice signals in somatosensory areas, even in non-tactile trials. Similar effects have been reported in an auditory task and persisted when correcting for movement-related activity^41^. Although subtle choice-predictive facial movements remain as a plausible explanation for this effect^70^, a closer investigation of choice formation in these somatosensory regions would be an interesting focus for future studies. Notably, such choice-related activity was absent in previous studies of a visual task in which mice reported their decisions by operating a steering wheel^20,71^, suggesting that it may be specific to directional-licking behavior. In this context, somatosensory choice signals might contribute to locating the lick spouts and supporting the execution of directional licks^72^.

Given the well-established role of MOs in choice formation in a variety of different tasks^22,39,43,54,73,74^ we focused on ALM to study single-cell activity during multisensory decision-making. Consistent with visual and tactile evidence being accumulated in RL and MM instead of ALM itself, we found no evidence of accumulated sensory signals. Instead, a large fraction of ALM cells represented the upcoming choices across sensory modalities. The temporal dynamics of choice representations closely matched the decoder and AUC results from widefield imaging, supporting a central role of ALM in cross-modal choice formation^22^.

In line with earlier work, we found roughly equal proportions of ipsi- and contralateral choice selective cells^73,75,76^. However, contralateral choice selectivity was stronger and emerged earlier in the stimulus period, suggesting that the contralateral choices were more directly driven by the sensory evidence, whereas ipsilateral selectivity may reflect interhemispheric transfer of choice signals. This could also explain the lack of correlation between visual and tactile choice AUCs in the late stimulus period when choice signals are still dependent on modality-specific sensory inputs. Only in the subsequent delay and response period, choice AUCs became modality-independent and highly correlated across visual, tactile, and multisensory trials, indicating a convergence toward a modality-unspecific choice representation. Notably, our recordings were restricted to layer 2/3 neurons while previous studies have shown that choice-related signals emerge earlier in deeper ALM layers^39^. Future studies could therefore aim to discern the layer-specificity of multisensory choice formation to determine if deeper ALM neurons show a stronger dependence on stimulus modality.

To causally test our imaging results, we inactivated either MOs or RL during different trial periods. We hypothesized that RL inactivation would primarily impair visual performance, while MOs inactivation would preferentially affect tactile performance and, critically, cross-modal choice formation. In visual trials, inactivation of RL and MOs impaired task performance in the stimulus and delay period, suggesting that both areas are involved in visual evidence accumulation, potentially due to bi-directional interactions^18^. In contrast, only MOs inactivation consistently impaired cross-modal task performance during the delay period, suggesting that multisensory evidence converges in MOs and is translated into behavioral output. The stronger impairment in tactile trials during the stimulus period further supports a role of MOs in tactile evidence accumulation, although imaging results pointed more specifically to area MM for tactile accumulation and ALM for choice formation^39^. Although our optogenetic inactivation was focused on ALM and we used low light power (64 mW/mm^2^) to achieve local effects, we cannot exclude a partial inactivation of the nearby MM^85^. The tactile bias with MOs inactivation might therefore arise from disrupting either ALM, MM or their inter-area interactions. Nevertheless, single-cell choice signals showed no modality preference and predicted choices equally during visual, tactile, and multisensory trials, arguing against a tactile bias in ALM choice formation.

Together, our work reveals how mesoscale cortical dynamics support multisensory decision-making, highlighting the convergence of unisensory evidence from parietal and frontal areas into cross-modal choice signals within MOs. Extending this framework to subcortical areas, such as the thalamus and superior colliculus will be important for establishing a brain-wide circuit view on multisensory decision-making. Our results also suggest that parallel multisensory pathways may exist for distinct behavioral demands. While we focused on temporal evidence accumulation and bilateral decision-making, future studies should therefore explore different task structures and ethological behaviors to investigate how these circuits either adapt to or are specifically tailored to diverse behavioral contexts.

## Methods

### Mice

All experiments were carried out in accordance with the German animal protection law and local ethics committee. Experiments were performed on 4 female and 10 male mice. At the time of surgery mice were between 8 and 12 weeks old. 4 female transgenic mice (2Niell/J^77^; JAX:024742 crossed with JAX:007004) contributed to the widefield-imaging data. 4 male C57BL/6J (JAX:000664), virally transfected to express the opsin stGtACR2 in CaMKIIα-positive neurons contributed to the optogenetics dataset. Additional 2 male mice (JAX:024742 crossed with JAX:007004) contributed to the two-photon dataset. The behavioral dataset consisted of both widefield- and optogenetics, as well as additional 4 male C57BL/6J mice only used in behavior. Mice were co-housed in groups of at least two animals and kept in a ventilated cabinet at a reverse day/night cycle (12/12 hours). Mice were provided with environmental enrichment and had access to food *ad libitum*. During behavioral experiments mice were water-restricted 5 days a week and provided with water as reward in the task. Water-restriction was interrupted on weekends.

### Surgical procedures

Surgical procedures were performed on mice aged 8-12 weeks. For widefield-imaging experiments, the skull was cleared using cyanoacrylate (Zap-A-Gap CA+, Pacer technology)^73,78^. For two photon preparations, a 3-mm-wide circular craniotomy was performed over the frontal cortex of mice, centered on the anatomical coordinates for ALM (2.5 mm anterior and 1.5 mm lateral from bregma). A glass coverslip was placed in the craniotomy and held in place by light-curable dental cement (DE Flowable composite, Nordenta) and fast curing acrylic resin (Jet Denture repair, Lang Dental INC). For optogenetics experiments, small craniotomies were made over the individual injection sites (frontal: 2.5 mm anterior and +-1.5 mm lateral from bregma; RL: 2.7 mm posterior and +-3.2 mm lateral from bregma) using a dental drill.

Viral injections were performed using a thin glass pipette with a micropump (UMP-3, World Precision Instruments) to deliver a viral vector (pAAV-CKIIa-stGtACR2-FusionRed; Addgene, viral titer ∼10^13^ vg/ml, diluted 1:10 in phosphate-buffered saline (PBS)) inducing expression of the inhibitory soma-targeted opsin stGtACR2^44^ in excitatory neurons. A total volume of 0.4 µl was injected per region. Subsequently, optical fibers (NA = 0.36, ø = 0.2 mm, FT200UMT, Thorlabs) housed in ceramic ferrules (ø = 1.25 mm, Thorlabs) were fixed to the skull above the injection sites using dental cement (Jet Denture repair, Lang Dental INC). Lastly, a custom stainless steel headbar was attached using quick adhesive cement (Superbond C&B, Sun Medical CO., LTD) and fast curing acrylic resin (Jet Denture repair, Lang Dental INC) allowing for subsequent head-fixation during the behavioral experiments.

### Behavioral task

After the initial learning phase, individual sessions of the behavior task consisted of interleaved visual, tactile and multisensory trials. Each of those stimulus conditions followed the same general schema (Figure 1a). A trial started with a 3 s stimulus period, in which mice were presented with sequences of sensory stimuli on their left and/or right side. Up to 6 sensory stimuli were presented at fixed time-points every 0.5 s throughout the stimulus period. In detection trials the maximum number of 6 stimuli were presented in absence of any distractors. In discrimination trials the presence of a stimulus in each time-bin was drawn randomly, with a probability of 0.7 on the target-side and 0.3 on the distractor-side, the same probabilities used in a previous study^38^. With this probability-based approach, there is a small chance of producing more stimuli on the intended distractor side. Therefore, target- and distractor side assignment and the associated rewards were defined based on the number of generated stimuli and not the underlying generative probability^38^. The stimulus period was followed by a 0.5 s delay to temporally separate sensation from the decision report and allowed for the study of evidence representation and choice formation. This period was followed by the 2 s response period, in which mice were presented with two water spouts on one on each side at which mice had to indicate their decisions in the form of licks to receive a small water reward (typically 2 to 3 µl). A response was counted once mice licked the same spout twice, consecutively. Individual trials were separated by an inter-trial-interval (ITI) of 1.5 s - 3.5 s. Mice were able to run on a custom 3D-printed wheel.

### Behavioral setup

Behavior setups were controlled using a microcontroller (Teensy 3.2, PJRC) running custom code in Python and C++. The microcontrollers (Teensy 3.2, PJRC) were used to control the stepper motors (setting the position of water spout; Nema 8, Pololu), solenoid valves (used for both water delivery and tactile stimuli; VDW22LA, 24 VDC, SMC), as well as photodiodes and lick sensors (capacitive detection). Behavior related data and triggers used for synchronization with physiological recordings were acquired using dedicated acquisition hardware (PCIe-6323, National Instruments).

### Visual stimuli

Visual stimuli were presented on two monitors (LG 23MB35PMF, 60 Hz refresh rate) mounted on the left and the right side of the mouse. Monitors were positions with their center at a distance of 18 cm from the eye of the mice. Covering ∼240° of the field of view in azimuth and up to 85° in elevation. Visual stimuli consisted of an individual cycle of a sine-wave with a spatial frequency of 0.018 cycles per degree (cpd). The magnitude was scaled by a factor of 1.25 and clipped to the range [-1, 1]. This effectively increased the contrast of visual stimuli. Stimuli were presented drifting from nasal to temporal at a temporal frequency of 2 Hz, with the black component of the stimulus leading (see Figure 1a).

### Tactile stimuli

Tactile stimuli were presented in the form of individual airpuffs, delivered via stainless steel tubes (16G gavage needle, straight, 125 mm long, AliExpress) mounted on top and in front of the mouse. Tubes were mounted at a distance of ∼15cm at an altitude of ∼35° from the horizontal plane and an azimuth angle of ∼10° directing air flow downwards and slightly away from the mouse. Spout positions were carefully calibrated to only deflect the distal part of the lower whiskers. Individual airpuffs were generated using short (20 ms) pulses triggering opto-coupled relays (AQZ105, Panasonic) switching solenoid valves (VDW22LA, 24 VDC, SMC). In turn, valve controlled the flow of compressed air (0.03 bar), resulting in a brief, subtle pulse of air. Tactile stimuli were synchronized to visual stimulus presentation using a photodiode signal.

### Behavioral training

Behavioral training was started on the detection conditions. In these conditions the maximum number of 6 stimuli are presented on the target-side in the absence of distractor stimuli. To not overwhelm the mice, they were initially trained only on visual detection trials until the mice reached stable performance of above 75% correct responses over three consecutive sessions. Once this criterion was reached, tactile detection trials were introduced with sessions now consisting of 50% visual and 50% tactile trials randomly interleaved. If mice did not reach a performance of above 75% correct responses within the first two sessions, visual trials were removed and sessions consisted of only tactile trials. Once mice reach reliable performance in tactile trials, visual trials were reintroduced if they had been removed before. After mice demonstrated that they reliably perform both visual and tactile trials, multisensory detection trials were introduced, each modality condition now comprising ∼33.3% of trials in a given session. At this stage the mice have learned the full multisensory task in the detection conditions.

In the final stage of training discrimination trials, consisting of a variable number of target- and distractor stimuli are introduced. The fraction of discrimination trials was gradually increased to 80% of trials in a session. With this training regime a high number (n = 12) of mice could be trained to perform this multisensory accumulation of evidence task.

### Bias-correction

To correct for side biases of mice a number of measures were taken. First, the water dispensing valves were carefully calibrated in regular intervals, ensuring precisely equal reward sizes. Second, the sequence of left and right target trials was generated in a pseudo-random manner. Generally, the probability of left-and right trials was equal. However, if by chance three consecutive trials were presented with the same target side, the probability of presenting the next target again on this side was reduced from 50%, and further to 25% from 4 consecutive trials onward. This was intended to break up longer blocks of trials with the same target side, that would biased animals to simply repeat previous responses. This condition was rarely reached and therefore only shifted the overall probability of switching instead of repeating target sides to ∼55%. Finally, to mitigate any remaining side-biases, the fraction of correct responses over the last ten left- and last ten right trials was compared and if strong imbalances were detected, the position of the preferred spout was automatically, gradually adjusted in small increments making the preferred spout slightly more cumbersome to reach until the proportion of left and right responses roughly equalized again. This primarily acted upon the initial learning phase of the task.

### Behavior videography

The movements of subjects were monitored using two CMOS-cameras (acA1920-155um, Basler) with the frame acquisition externally triggered (90 Hz) to achieve synchronization with the widefield imaging. One camera captured facial movements and one overall body movements. Video data was compressed using H.264 encoding with a compression factor of 17, implemented in python using “ffmpeg” (Tomar, 2006^79^).

### Widefield imaging

The widefield Ca^2+^-imaging was based on an inverted tandem-lens macroscope^80^, with a 50 mm objective (Nikon AF-D 50mm f1,4; Foto Erhardt) above the subjects and a 85 mm objective in front of a sCMOS camera (Edge 5.5, PCO). A schematic of this macroscope is shown in Figure 2a. Frames were acquired using custom software written in LabView (National Instruments) at an imaging rate of 30 Hz using 4 x 4 spatial binning resulting in a resolution of 640 x 540 pixels. Frames were alternatingly illuminated using a blue LED (470 nm, M470L3, Thorlabs) and a violet LED (405 nm, M405L3, Thorlabs) with a 405 nm excitation filter (#65-133, Edmund optics). LEDs were collimated using adjustable collimator lenses (SM2E, Thorlabs). LEDs were powered using LED-drivers (Cyclops LED Driver 3.6, Open ephys). Both excitation light paths were merged using a dichroic mirror (no. 87–063, Edmund optics) and reflected onto the brain using a second dichroic mirror (G381323036, Qioptiq). GCaMP fluorescence signals were acquired using a 525 nm emission filter (MF525-39, Thorlabs) mounted in front of the camera. By alternatingly illuminating frames using either the blue or violet LED, two separate movies at 15 Hz each are acquired. Here, frames acquired under blue illumination reflect Ca^2+^-dependent fluorescence, while frames acquired under violet illumination represent Ca^2+^-independent fluorescence signal^74,81^. This allowed us to compensate for Ca^2+^-independent signal by subtracting the rescaled frames acquired under violet illumination, from the preceding blue illuminated frames. All analysis was based on this differential signal.

As part of the preprocessing, widefield imaging data was motion corrected using a subpixel image registration routine^82^. This was performed separately for blue- and violet movies before computing the differential calcium-dependent signal. To facilitate the analysis of widefield data, we used SVD (singular value decompositions) to compute the 300 highest variance dimensions of the data. This results in ‘spatial components’ U (of size pixels x components), ‘temporal components’ V (of size components x frames) and singular values S (of size components x components) scaling components to match the original data. All analysis is performed on the scaled temporal components SV. SV was high-pass filtered above 0.01 Hz using a zero-phase, second-order Butterworth filter. Following analysis of SV, signals were projected back into the original pixel space by convolving with U. All widefield data were rigidly aligned to the Allen CCF^83^, using anatomical landmarks: left, center and right points where anterior cortex meets the olfactory bulbs, and central point at the base of retrosplenial cortices. This alignment was functionally confirmed using visual field sign mapping^84^ (Figure 2a).

### Two-photon microscopy

Two-photon imaging was performed with a custom build resonant-scanning (CRS 8KHz, Cambridge Technology) microscope with a Ti:Sapphire femtosecond pulsed laser (Mai-Tai DeepSee, Spectra-Physics) and a 16x 0.8 numerical aperture objective (Nikon Instruments). The microscope was controlled using the Matlab-based package Scanimage^85^. Fluorescence Images were acquired at 30.4 Hz with a resolution of 512 x 512 pixels (575 μm × 575 μm) using an excitation wavelength of 920 nm. Recordings were made from a single plane in layer 2/3 of MOs at a depth of 250-300 µm below the pial surface. During recording, small shifts in the z-plane were manually corrected when needed.

Two-photon imaging data were processed using the Suite2P package^86^, to perform rigid motion correction, model-based region of interest (ROI) detection, correction of neuropil contamination and spike inference. Somatic ROI identification was performed through a combination of a pre-trained classifier and manual curation. ΔF/F traces for each neuron were then produced using the method of Jia et al.^87^ but skipping the final filtering step.

### Optogenetic inactivation

Photostimulation was performed using a 473-nm Laser (LuxX 473-100, Omicron). With a power density of 64 mW/mm^2^. Stimuli consisted of a brief ramp up of 100 ms, a sustained plateau and a longer ramp down over 500 ms, to avoid an excitatory post-illumination rebound due to sudden release from inhibition ^88^. To prevent animals’ from visual detecting the photostimulation, a removable light-shielding was attached to the head-fixation during behavioral sessions. Once mice had learned all three detection conditions, bilateral optogenetics inhibition trials were introduced (Figure 7d). Photostimulation trials were randomly interleaved with control trials, while prohibiting two consecutive photostimulation trials. For ’Stimulus’ inactivations the ramp up began 500 ms before the start of the stimulus period and the ramp down was finalized at the end of the stimulus period. For ‘Choice’ inactivations the ramp up instead started 100 ms before the end of the stimulus period to ensure full inactivation during the delay and the ramp down only began at the end of the response period.

## Data Analysis

### Behavioral performance

The performance of mice was calculated as the fraction of correct responses over all responded trials. Learning curves were computed as the performance over sessions since a modality condition was introduced to a given mouse (Figure 1b). For this we determined the relative offset in the number of sessions when the tactile and multisensory conditions were first presented to an individual. This was used to compare performance in the tactile and multisensory condition relative to the number of sessions a mouse had performed this condition. Here, we compared the performance between visual and tactile and between tactile and multisensory conditions for individual sessions since introduction across mice using Wilcoxen-Man-Whitney U tests.

Next, to compare if mice integrate visual and tactile stimuli to improve their performance, we compared the performance in multisensory trials to the individually preferred unisensory modality. We fit the following LME model to this data using MATLAB’s fitlme function:

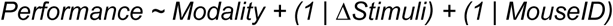

Here, Performance denotes the fraction of correct responses, Modality is the fixed effect of either the multisensory or the preferred unisensory condition and the random effects ΔStimuli as the difference in target and distractor stimuli and MouseID affecting the intercept. The psychometric curves presented in Figure 1e, were obtained using the Matlab-function ‘fitbnd’ fitting the sigmoid function:

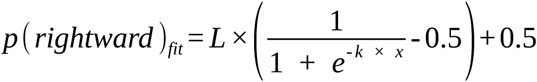

Here, L defines the maximum performance (limited to [0, 1]), k scales the slope of the psychometric curve and x denotes the difference in right- and left sensory stimuli. For the presentation of performance as log odds ratios, the ratio of individual probabilities of rightward- and leftward responses was computed and presented on a logarithmic scale (Figure 1e, bottom; y-axis). The log odds ratio was computed as:

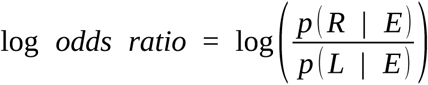

Here, p(R|E) and p(L|E) represent the probabilities of responding right- or leftwards given the sensory evidence presented. E represents the sensory evidence expressed as the number of excess stimuli presented on the right-side. Accordingly, evidence represented right-target trials as positive integers and left-target trials as negative integer values. We then tested if the performance in multisensory trials could be described as the additive integration of visual and tactile evidence, applying the relationship presented by Coen et al.^22^:

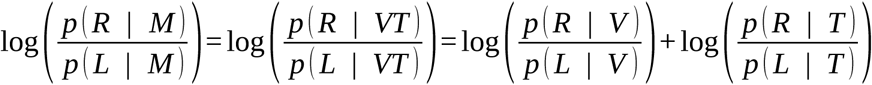

Here, M represents multisensory evidence and VT indicates the additive combination of visual and tactile evidence, under the assumption that this is equivalent to M. V and T represent visual and tactile evidence, respectively. Note that we removed the bias-term in this equation, as the experimental design of the behavioral setup actively corrected for side-biases. We modeled the log odds ratios for the individual modality conditions using a linear regression. For this, we computed the slope of the linear fit as:

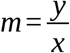

Here, x denotes the sensory evidence as the difference in right- and left stimuli, y represents the log odds ratios and m represents the slope of the linear transformation describing how strongly the difference in the number of sensory stimuli affects the performance of mice (as log odds ratios). By expressing the performance of mice in this linearized form we were able to make a prediction for the optimal additive integration of visual and tactile sensory information assuming:

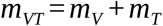

Here, m_VT_ denotes the theoretical slope of the linear transformation under the assumption of an optimal additive integration of visual and tactile information. m_V_ and m_T_ denote the computed slopes obtained from performance in unisensory visual and tactile trials, respectively. This allowed us to compare the similarity m_M_ to of this theoretical m_VT_, as well as to to performance in the individually preferred unisensory condition. The additive integration similarity was computed as:

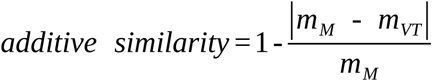

And the similarity to the fit of the preferred unisensor modality performance:

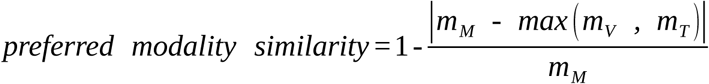

By comparing “additive similarity” with “preferred modality similarity” using a two-sided signrank test, we tested if the performance in multisensory trials was better described by the assumed additive integration of visual and tactile information compared to the performance in the individually preferred unisensory condition (Figure 1g).

### Psychophysical Reverse correlation analysis

The reverse correlation analysis used in this study is based on the approach presented by Scott et al.^38^. With the difference that we compared the empirical excess-stimulus probabilities not to the theoretical difference, but the empirically determining difference in excess stimuli in the respective time-bins. This analysis reveals to which extent sensory stimuli in a certain time-bin influenced the behavioral choices of animals. Here, we computed the choice conditioned excess-stimulus probabilities depending on the relative time of presentation within the stimulus period of the task. This analysis was performed independently for visual, tactile and multisensory discrimination trials using student’s t-tests, (n=12 mice; Figure 1h). The additive integration prediction in Figure 1i, was computed as the mean probabilities of excess stimuli across visual and tactile trials (as multisensory stimuli were assumed to represent an individual, more complex stimulus, instead of two cross modal stimuli) and compared these to the observed stimulus weights computed from the performance in multisensory trials. For this comparison, used the following LME model:

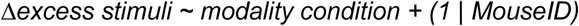

Here, Δexcess stimuli is the difference in excess stimuli between multisensory trials and the additive prediction. Modality condition denotes these two modality conditions as a fixed effect and MouseID is the random effect on the intercept.

### Detection trial responses

Maps and traces of correctly responded-detection trial activity (Figure 2b-d, Supplementary Figures S2-4) were generated by averaging responses across mice, with 25 sessions each. Here, we first computed session-wise average responses that were subsequently combined using a weight mean based on the number of trials contributing in each session. Activity was expressed as ΔF/F by referencing activity to a 1s baseline before the stimulus onset. We combined left- and right target-trials, by reflecting the hemispheres in left-target trials so maps and traces instead represented ipsi- and contralateral trial activity. Spatial maps of activity were generated by averaging activity of defined time-bins (usually 0.5s). To obtain region-wise traces of activity over time, activity was computed for individual regions of interest (ROIs) using a circular ROIs with a diameter of ∼750 μm. Traces are presented as mean across sessions and mice. Next, we compared the responses in multisensory trials to the unisensory conditions using a linear-mixed effects model based on the average response during the stimulus period (Figure 2c,d):

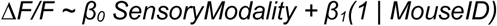

Further, we determined the overlap in activity across visual and tactile trials, expressed as the unisensory overlap index (Supplementary Figure S3). For this, we expressed activity in units of standard deviation across sessions and mice. Next, we computed the product of the normalized responses in visual and tactile trials and termed this the unisensory overlap index as this metric represents strong and reliable responses found across both unisensory trial conditions. Next, we compared activity in multisensory trials to either the higher unisensory response (Multi. - max(Uni.)) or the summed activity across both unisensory conditions (Multi. - sum(Uni.); Supplementary Figure S4). These operations were performed for each pixel individually after computing the average activity across the time bin.

### Linear encoding model

We used a linear encoding model to test if sensory responses in multisensory trials could be fully described by an additive integration of visual and tactile sensory responses. For this purpose, we designed a linear encoding model that predicts the recorded neural activity based on task- and movement related activity. The linear encoding model used in this study was based on a previously published model design^40^.

The model consists of individual task- or behavior related signal regressors combined into a single design matrix used to describe the recorded neural activity (Figure 3a). One the one hand, this design matrix was composed of analog regressors including running speed, video regressors and the video motion energy (ME). On the other hand, sets of binary regressors each represent a specific time-point (frame) of the same related event, but shifted in time. This allowed them to model time-varying event-related responses.

To model responses to the sequences of sensory stimuli, we distinguished between the response to the first sensory stimulus at the onset of the sequences and responses to the subsequent stimuli. The first stimulus response was modeled using time-varying regressors from the onset of the first stimulus to the end of the delay period. Subsequent sensory stimulus responses were modeled from 0.5 s before to 1 s following the onset of each subsequent stimulus.

With this combination of first and subsequent stimulus regressors, stimulus sequences on the left- and right side were modeled independently. Visual and tactile stimulus responses were independently modeled in both unisensory and multisensory trials condition. In multisensory trials both the visual and tactile stimulus regressors were active simultaneously to describe additive multisensory responses. In addition, a third type of sensory regressor with the same characteristics as the visual- and tactile regressors was used to describe non-linear multisensory responses, whenever visual and tactile stimuli were presented simultaneously (Figure 3a). These non-linear multisensory regressors were designed to capture stimulus related responses that deviate from the additive integration of visual and tactile stimuli. Other task-related signals, such as trial time, choices, outcome as well as previous trial choice and outcome were modeled using regressors covering the entire trial duration.

The model also contained information about movements of the animals. Movement-related responses were modeled using event-related regressors from 0.5 s before to 1.5 s after the respective events. These included individual left- and right licks as well as the onsets and offsets of running. Furthermore, the model contained analog regressors, including the running speed resampled to the imaging rate as well as the 200 dimensions with the highest variance representing the behavioral videos obtained using SVD (similar to Widefield imaging). In addition, the motion energy (ME) computed individually for the two behavioral cameras was included. Supplementary Table 1 provides an overview of all model regressors.

To be able to include video regressors, we had to ensure that these didn’t contain redundant information already described by other regressors in the model, such as the sensory stimuli, licks and running. For this purpose, the design matrix was arranged with video- and ME regressor columns at the end of the matrix. We then performed a QR decomposition^89^ and replaced the original video- and ME regressors with the new orthogonalized regressors^40^. This was critical to ensure that no explanatory power about the sensory stimuli (visual in particular) could be obtained from the videography data.

We used an analysis of the cumulative subspace angles^40^ to ensure that individual regressors in the design matrix were not fully redundant, which otherwise would result in rank-deficiency of the design matrix. The model was fit, using ridge regression, to avoid overfitting. We estimated the regularization penalty separately for each column of the widefield imaging data using marginal maximum likelihood estimation ^90^. Minor modifications were made to reduce the numerical instability for large regularization parameters. We used this model to isolate sensory responses from other task- and behavior related activities and tested if sensory responses could be fully described as an additive multisensory integration, or if a non-linear integration term could improve the model fit.

### Variance analysis

To estimate the relative contributions of sensory- and movement-related information to the model predictions, we computed the explained variance (cvR^2^) using tenfold cross-validation (Figure 3b). To determine the unique explained variance (ΔR^2^) we generated reduced models, identical to the full model, with the exception that selected regressor groups were shuffled in time to destroy their correlation with the widefield data. To account for the effect of shuffling itself (particularly for the interpretation of multisensory regressors) the full model contained a full copy of the multisensory regressors, which were already shuffled in the full model. Reshuffling these regressors served as control for the unique explained variance analysis. We computed the cvR^2^ for all reduced models and compared these to the full model to obtain the ΔR^2^ (Figure 3c). Next we compared the ΔR^2^ of each reduced model to the shuffle control using an LME.

### Sensory response kernels

To inspect the response kernels used by the linear model to describe the sensory stimulus response, we compiled the β-weight kernels for the individual first sensory stimulus regressors across sessions and presented the average weight over the first 1 s by convolving with U. For regions of interest with high kernel weights, we computed average kernel traces by averaging over pixels in the reconstructed original pixel space, on the level of individual sessions and present these as mean ± s.e.m. across mice (Figure 3d).

### Unisensory trained model

As an alternative approach to determine the extent to which information about sensory responses in unisensory trials would suffice in predicting multisensory trial activity, we conceived an alternative model fitting approach. Here, we use the same tenfold cross-validation procedure previously described, but fitting the full model either on only the unisensory trials of a session (termed uniModel) or fitting the same model on a matching number of trials but containing unisensory as well as multisensory trials (termed matchedModel). We evaluated these model predictions on the subset of multisensory trials (∼33% of multisensory trials per session) that were excluded from the cross-validated fit of both models, by comparing these predictions to the measured cortical activity. We present these model predictions either as traces for selected regions of interest, or as the summed absolute deviation from the measured activity over the entire trial (Figure 3e). We compared the uniModel as well as the matchedModel to the measured cortical activity using the following LME model:

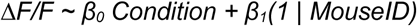

Here, Condition denotes either the measured activity or the respective model prediction and MouseID the identity of the animal.

### Sensory evidence- and choice-related activity

To distinguish between activity related to the representation of accumulated sensory evidence (Figure 4, Supplementary Figure S5) and behavioral choice (Figure 5, Supplementary Figure S6), we needed to decorrelated stimulus- and choice related activity, which would otherwise arise from mice performing the discrimination task at higher than chance level. We use a trial balancing procedure, previously presented^39^, where we equalize the number of correct and incorrect, left- and right trials, so that sensory information was no longer related to the decision of mice. Then, we computed the difference between activity of the left- and right hemisphere to determine which cortical regions reliably represented sensory evidence- or choice-related information by computing the area under the receiver-operating characteristic curve (AUC). For this, we used a bootstrapping approach to generate 100 random samples of trials performed by an animal across multiple sessions. Here, we randomly selected 40 choice-balanced discrimination trials out of all trials performed and computed the AUC for either the sensory evidence or the upcoming choices. We compared the sensory- and choice-related AUCs across ROIs during the delay period of the task using LME models:

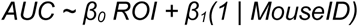

Here, ROI denotes the activity of activity of pairs of ROIs we compared and MouseID the identity of the animal.

### Neurometric curves

We compared how reliable RL and MM represented the target side in discrimination trials of varying task difficulties. For this, we used the same bootstrapping approach described in the previous section, selecting trials of either easy (ΔStimuli = [3,4]) or hard (ΔStiuli = [1,2]) task difficulty. For each, we computed the AUC during the delay period by sensory modality. The resulting neurometric curves are then presented as mean ± s.e.m. across mice. To compare AUCs across conditions, we used the following LME model:

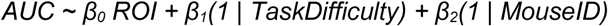

Here, ROI denotes the activity of pairs of ROIs we compared, TaskDifficulty the difficulty levels of trials as either easy or hard and MouseID the identity of the animal recorded from.

### Choice decoder analysis

To determine the time-course of choice formation in this task, we used a previously presented decoder approach^41^. This decoder was designed to predict the animal’s left/right choices based on the widefield imaging data, combining signals across the entire dorsal surface of the brain. We trained this logistic regression decoder with an L1 penalty on the temporal component matrix SVT for individual sessions. The L1 penalty was defined as the inverse of the number of observations in the test dataset during cross-validation, which yielded a good balance between the cross-validated prediction accuracy of the decoder and the number of nonzero model regressors.

We matched the number of correct and incorrect, left- and right choice trials, by determining the trial combination with the lowest trial count and randomly selecting the matching number of trials from all other combinations (at least 60 trials). This procedure ensured that sensory responses or side biases could not be used by the decoder to predict choices. To reduce overfitting we reduced the dimensionality of the imaging data to the 50 dimensions with the highest explained variance. The logistic regression model was implemented in MATLAB using the ‘fitclinear’ function and run individually for each time point within a trial. We computed the decoder prediction accuracy using tenfold cross-validation at each time point individually. β-weight maps were computed by averaging over all models fit during the cross-validation and projected these back into the original pixel-space by convolving with the spatial components U.

In order to best utilize all available data, we used the maximum number of balanced trials within a session. We accounted for the variable number of trials between sessions by computing mouse-wise weight averaged for the prediction accuracy and β-weight maps.

### ALM cell selectivity

We characterized the stimulus and choice representation of ALM neurons obtained using two-photon microscopy (Figure 6). With the exception of the visualization in Figure 6b, analysis was performed on the inferred spikes, computed using suite2p. Activity of cells was referenced to the 1s baseline preceding the stimulus onset across all trials within a session. First, we determined the fraction of cells that were responsive to sensory stimuli of either modality condition. Here, we averaged activity over the entire 3s stimulus period of correctly-responded detection trials. We compared this activity to the activity during the baseline period. To ensure that sparse activity does not bias this analysis, we pooled activity of 3 randomly selected 1s baselines throughout the session. We determined the fraction of cells that significantly responded to the sensory stimuli using a one-sided signrank test (threshold p<0.05). Using a binomial test, we compared the fraction of significantly responding cells to a control in which the identity of baseline and stimulus was shuffled (Figure 6c, left). Next we determined if ALM cells reliably represented either the target-side or the upcoming choices in choice-balanced discrimination trials. We determined the fraction of significantly responding cells by comparing the activity during the 1s delay period (extended for two-photon mice) between ipsi- and contralateral target/choice trials using a ranksum test. These were then compared to a shuffle control using a binomial test (Figure 6c, middle and right). These metrics were computed for each modality condition separately. To determine the time course of choice formation in the ALM neural population, we used the inferred spikes averaged across a moving window of 1s over the course of trials. We denoted the end of this moving window as the time point of the response, reflecting activity over the preceding 1s of the trial. We computed the fraction of choice selective cells as previously described, now across time. We compared these fractions of cells to a shuffle control using a binomial test and determined the onset of choice selectivity for each modality condition (Figure 6d).

To compare the representation of choices, we computed AUCs for each cell, reflecting how reliable these represent either ispi- or contralateral choices. For all cells that were choice selective for at least 1s in one of the modality conditions, we presented the average choice-related AUC (Figure 6e). For this, cells were categorized as either ipsi- or contralaterally choice selective depending on the distance of the lowest and highest 2.5 percentile of AUC values from 0.5. Here, AUC values below 0.5 indicate increased activity in ipsilateral choice trials and values above 0.5 indicate increased activity in contralateral choice trials.

To determine if the same cells represent choices across modality conditions, we compared the choice-related AUCs between conditions for the last 1s of the stimulus period, the delay period and the first 1s of the response period. Here, we compared AUCs between visual and tactile trials, as well as comparing AUCs in multisensory trials to the higher (furthest from 0.5) AUC across both unisensory conditions using Spearman correlation.

### Optogenetics

We determined the causal effects of bilateral optogenetic inactivations of RL and MOs. For this, the control performance of mice was determined in the individual modality conditions across detection and discrimination trials. Then performance in trials with either ‘Stimulus’ or ‘Choice’ inhibition was compared to this control performance using a binomial test. Confidence intervals were computes as the Wilson score intervals.

## Acknowledgements

We thank Madgalena Robacha for establishing preparations; Alex Bexter for his work and gained insights on the preliminary version of this visuotactile behavior task; Mira Ritter, Lena Kricsfalussy-Hrabár, Sandra Brill, and Severin Graff for assistance with behavioural training; Elisabeta Balla and Thomas Rüland for assistance with the two-photon experiments and data preprocessing and Michael Moll for providing the computational infrastructure. This work was supported by the Deutsche Forschungsgemeinschaft (DFG, German Research Foundation) - 368482240/GRK2416 and 430156848/SPP2205, the Helmholtz association (VH-NG-1611) and the state of North Rhine-Westphalia through the iBehave initiative. Furthermore, this work was funded under the Excellence Strategy of the Federal Government and the Länder.

## Author contributions

B.M.K., S.M. and G.N., conceived and planned the experiments. G.N. designed the behavioral task and engineered setups and microscopes used, collected behavioral- and neural data and performed the analysis with input from B.M.K. and S.M.. G.N., I.L., S.A.R., E.C., conducted the behavioral training of animals and collected the neural data. G.N. wrote the initial draft and generated the figures with input from B.M.K and S.M. The manuscript was revised and completed by B.M.K and S.M. Correspondence should be directed to Björn Kampa, and Simon Musall.

## Supplemental information

**Table 1:**
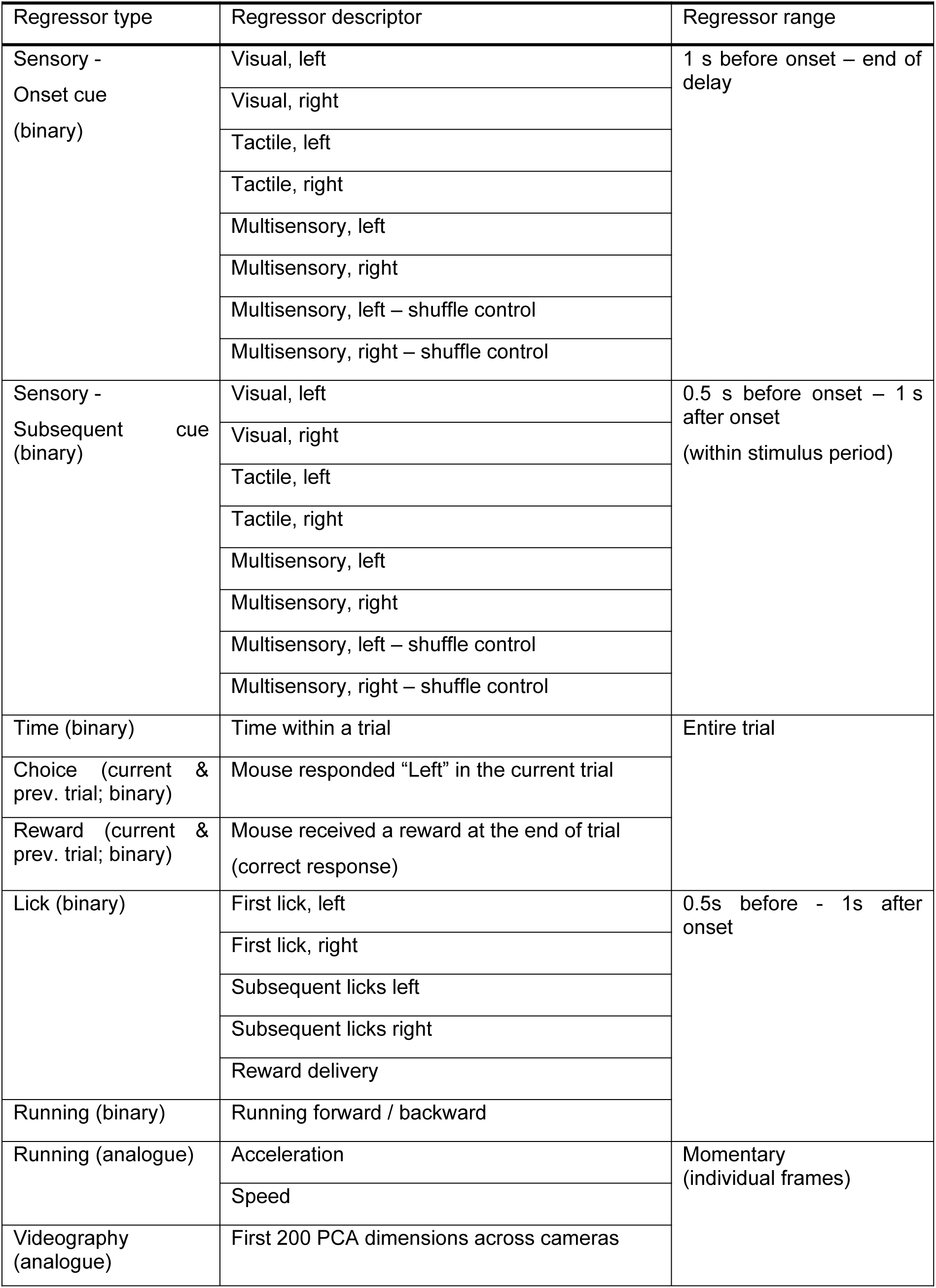
Linear model regressors.

**Figure S1:**
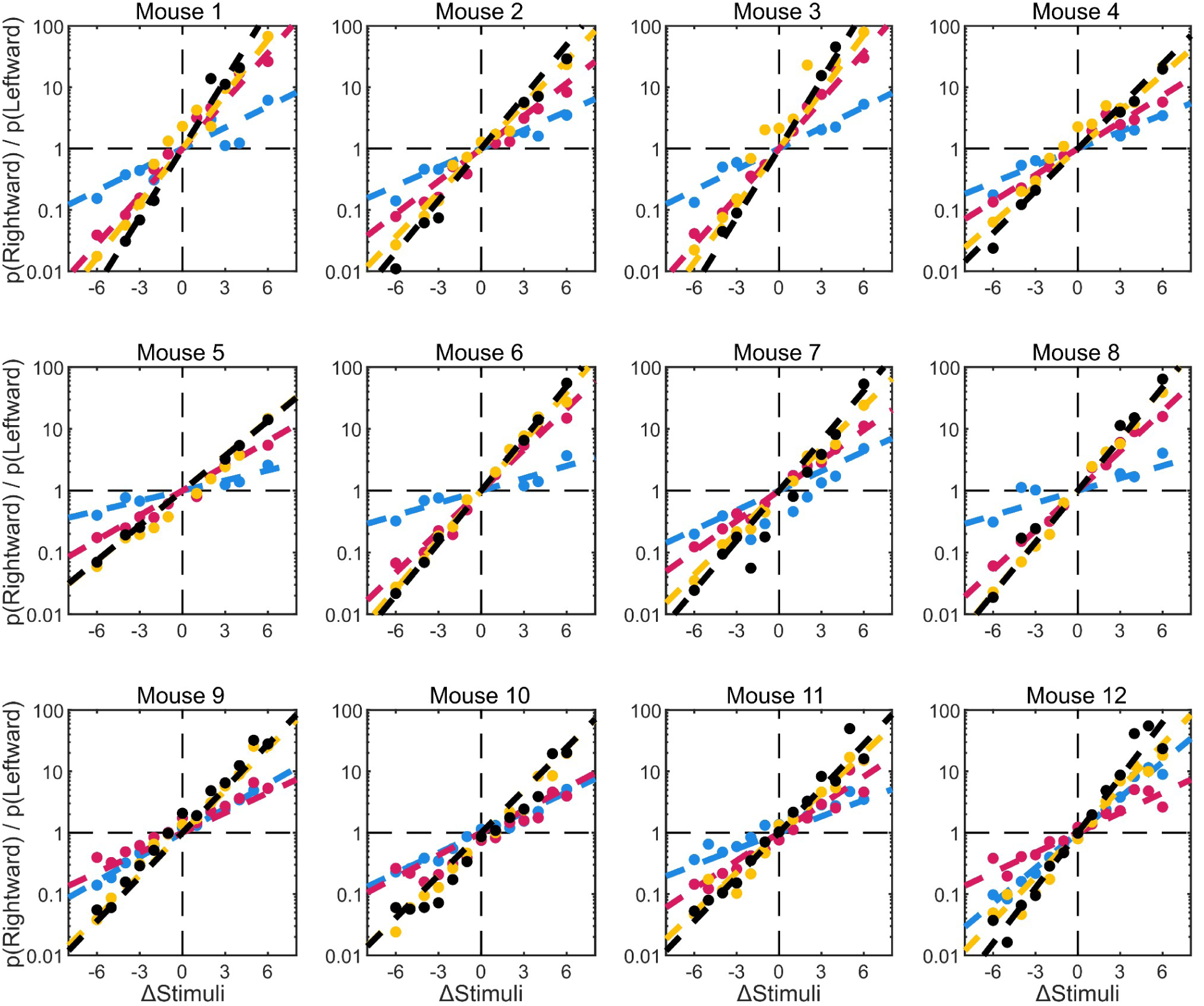
Multisensory performance of individual animals is well predicted by the additive integration. Task performance, presented as of log-odds ratios for individual mice, analogous to Figure 1f.

**Figure S2:**
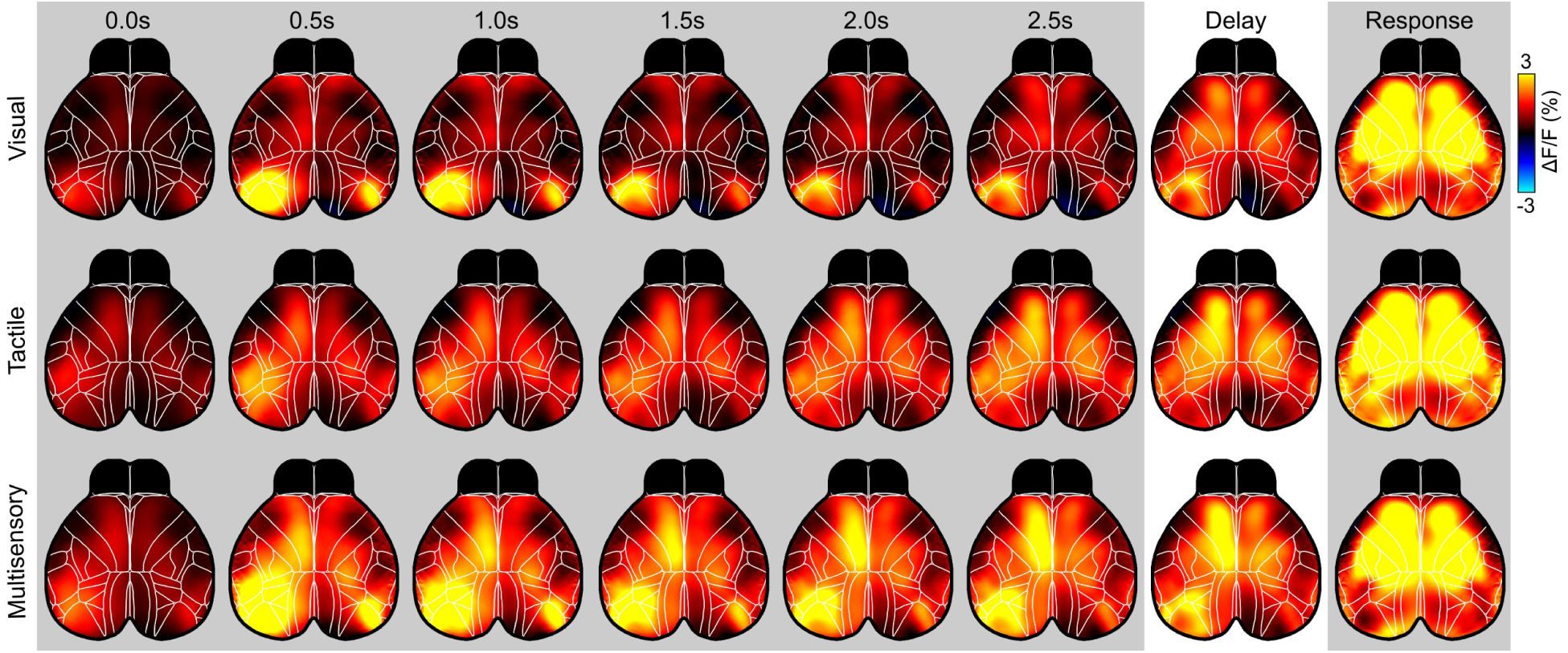
Average detection trial activity. Average cortical activity of mice performing the three stimulus conditions. Activity is shown for rewarded detection trials with the maximum number of target stimuli without any non-target stimuli (n = 100 sessions from 4 mice). Left- and right trials were combined by reflecting data from left-target trials along the midline to represent contra- and ipsilateral activity relative to the target side instead of the left and right hemispheres, respectively. Column labels indicate the start of each 0.5s time-bin. Correspondingly, the bin labeled 3.0s denotes the delay period and bin 3.5s the response period. Mice displayed similar patterns of cortical activity throughout the stimulus period. Towards the end of the stimulus period over the delay and response period, activity became more similar with increasing activity in the frontal cortex across modality conditions.

**Figure S3:**
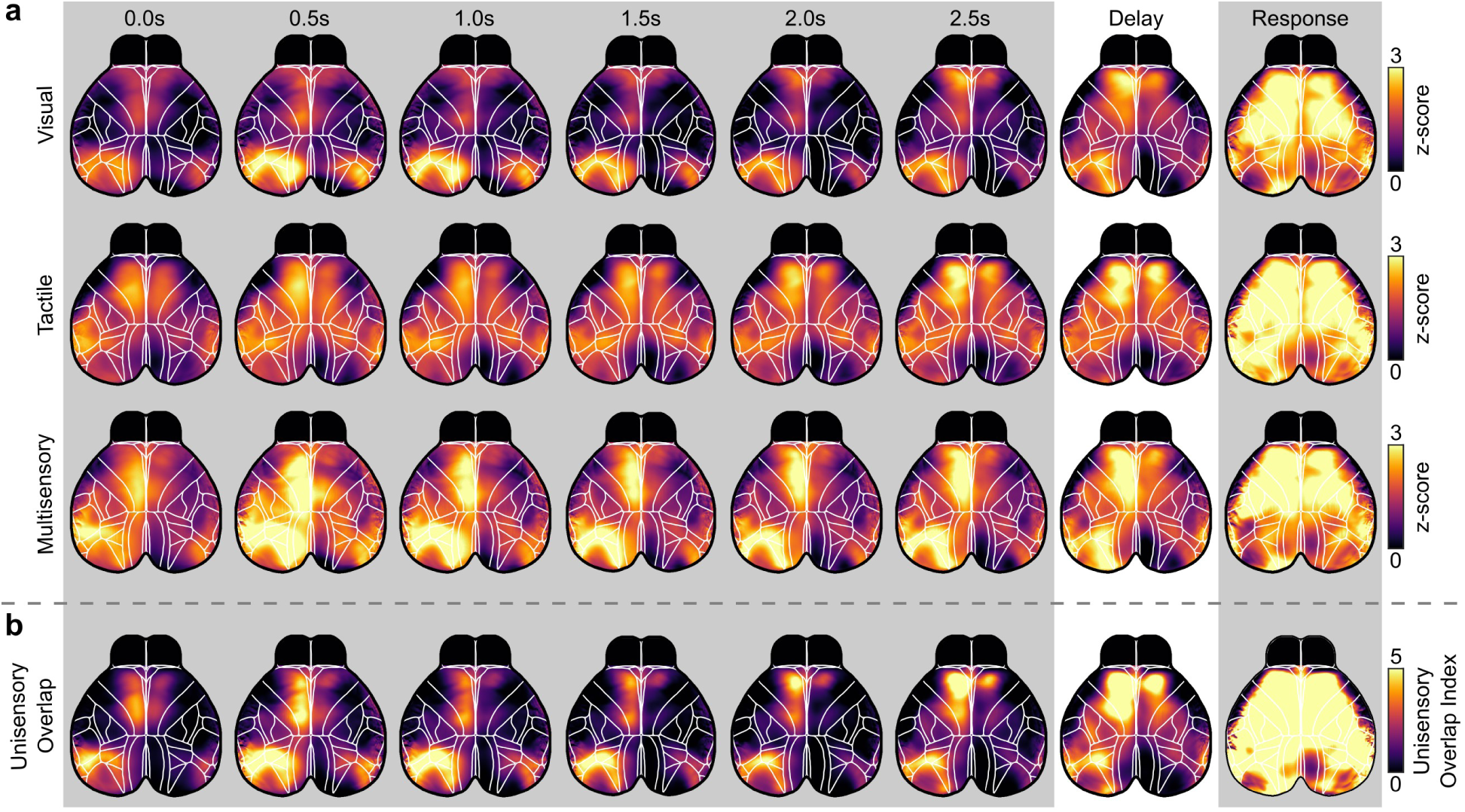
Overlap of unisensory trial activity in RL and MOs. (**a**) Same maps of average cortical activity in detection trials, presented in Fig. S2., here expressed in units of standard deviation. (**b**) Unisensory overlap index computed as the product of the normalized responses in visual and tactile trials shown in (a). Extended presentation of maps displayed in Fig. 2e. Throughout the stimulus and delay period, the largest overlap in activity was found in the HVA, particularly area RL and the MOs. Here, RL displayed the highest overlap index early in the stimulus period, while this over increased towards the end of stimulus and during the delay period for MOs.

**Figure S4:**
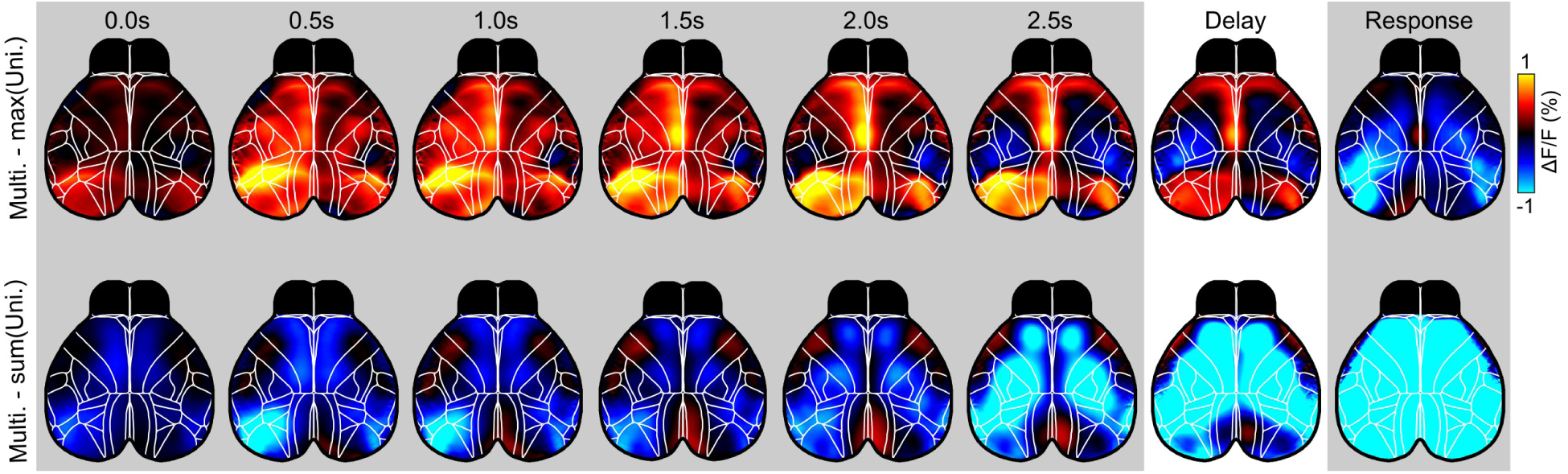
Multisensory trial activity is higher compared to both unisensory trials. Top: Comparison of multisensory trials activity to the unisensory trials (Multi. - max(Uni.)). Here, we compared the activity of each pixel to the higher activity across visual and tactile trials. Presented for 0.5s time bins over the course of correctly responded detection trials shown in Fig. S2. In particular, RL and MM display higher activity in multisensory compared to unisensory trials. Bottom: Comparison of multisensory trial activity to the summed activity across both unisensory conditions (Multi. - sum(Uni.)). Throughout cortex, activity in multisensory trials tended to be lower compared to the summed unisensory trial activity.

**Figure S5.**
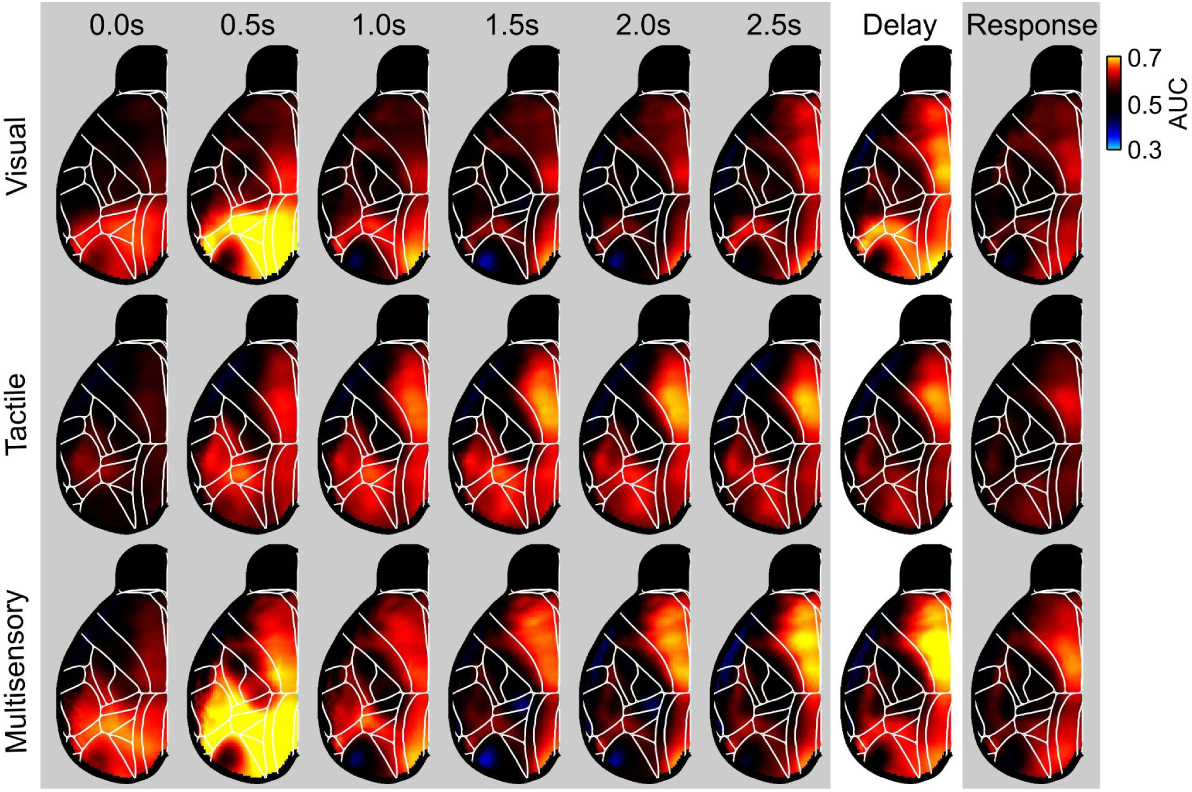
Modality-specific sensory evidence accumulation across trial time. Representation of sensory evidence in different stimulus conditions. Analogous to Fig. 4a, but presented for all 0.5s time bins from the onset of the stimulus to the initial response period. Presentation The AUC maps show cortical regions that reliably represented the side where the higher number of sensory stimuli was presented, based on the difference in activity between hemispheres. AUCs were computed for discrimination trials with an equal number of correct and incorrect trials to avoid an influence of choice-related activity. AUCs above 0.5 indicate a more reliable representation of sensory evidence.

**Figure S6.**
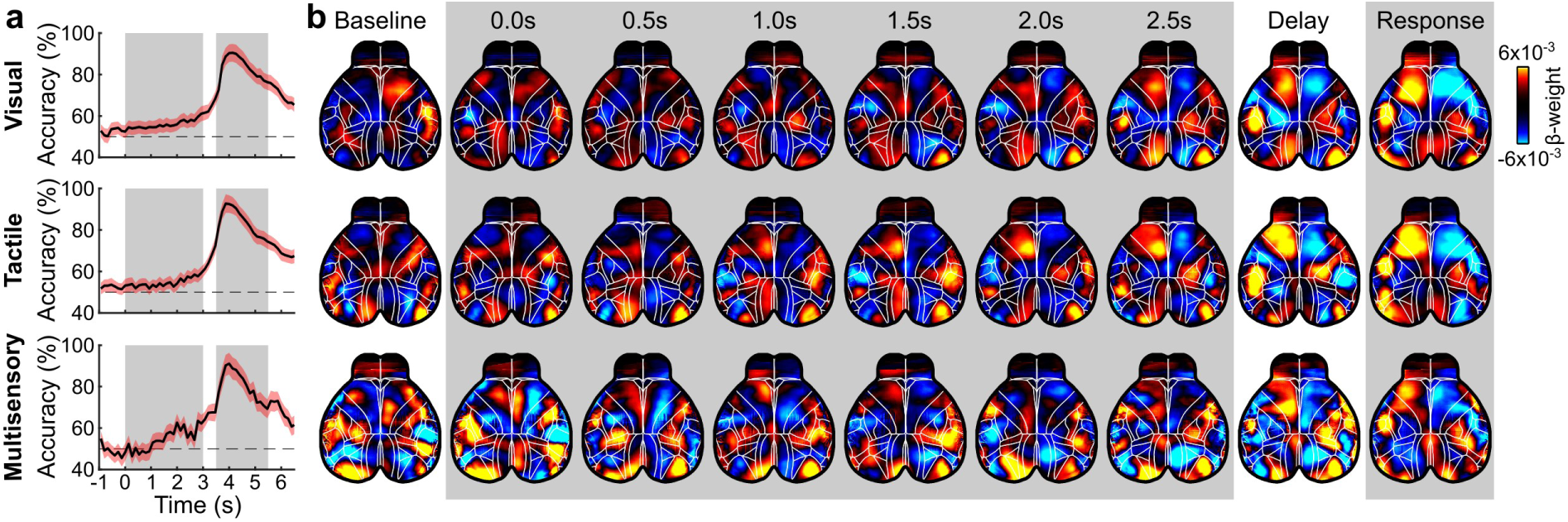
Modality-specific choice decoders. Individual choice decoders (matching Figure 5a,b) were fit to either visual, tactile or multisensory trials. (**a**) Prediction accuracy of each decoder over time, presented as mean (black) and s.e.m. (red shading) across mice. (**b**) Beta-weight kernels across time. Average beta-weight over 0.5s time bins shown for baseline, stimulus, delay and response.

**Figure S7.**
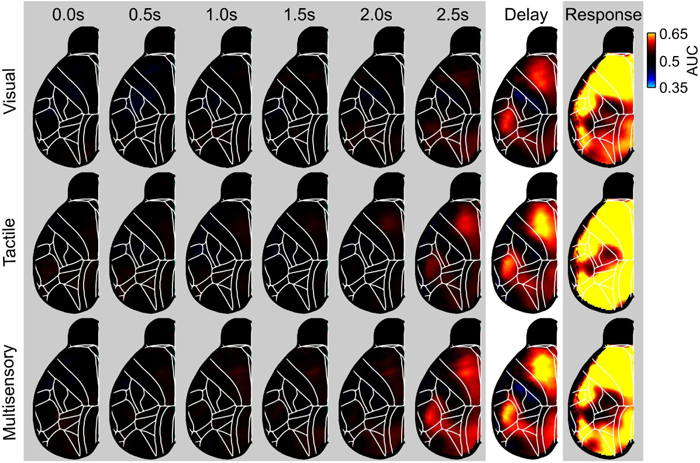
Modality-unspecific choice formation across trial time. Spatial maps of AUCs reflecting upcoming choices in the different stimulus conditions. Analogous to Fig. 5c, presenting the full time-course from the stimulus onset to the initial response period in time bins of 0.5s.

**Figure S8.**
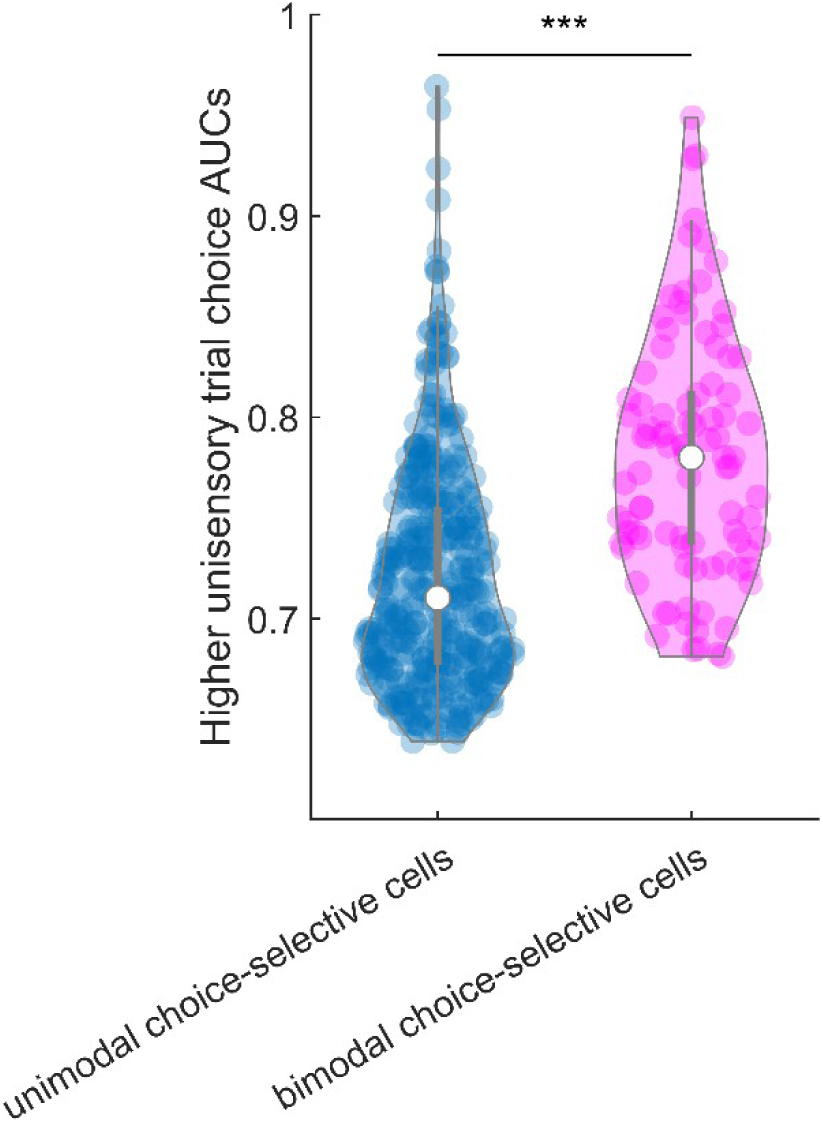
Bimodal choice-selective ALM cells more reliably represent choices. Choice-related AUCs during the delay period of the task (data presented in Figure 3i). Cells were grouped based on if they significantly represent choices exclusively in one of the two unisensory trial conditions (visual, tactile; denoted unimodal choice-selective cells), or if they reliably represented choices in both visual and tactile trials (bimodal choice-selective cells). The higher AUC-value across visual and tactile trials was determined and compared suing a ranksum test (p=1.6 x 10^−16^, n=453 cells).

**Figure S9.**
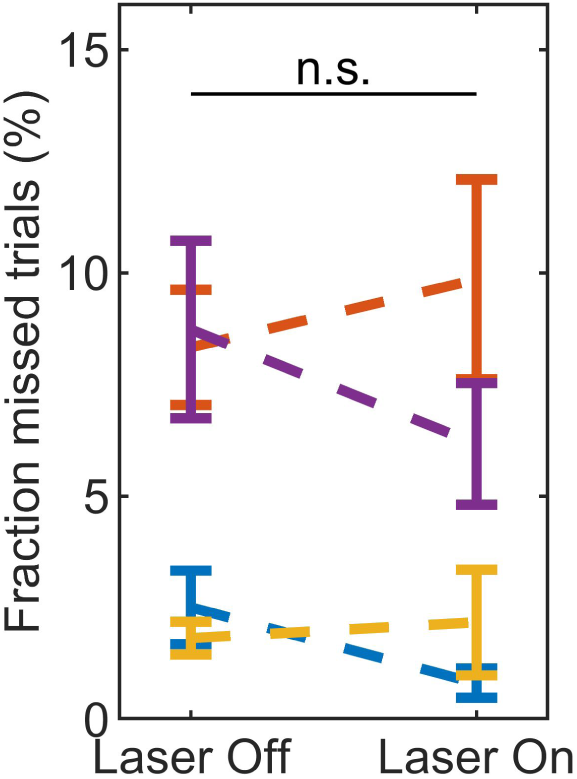
ALM inactivation during delay- and response period does not reduce lick-responses. As ALM is a region of MOs, one could imagine that inactivation affects the expression of the report behavior. To test this possibility, we computed the fraction of missed trials (trials without lick-resonses), comparing Laser Off and Laser On condition in sessions with ALM inhibition during the choice period of the task (delay and response). Presented for individual mice as mean ± s.e.m. Compared using a LME model (p=0.48, T=-0.71, LME model, n=144 sessions from 4 mice).

